# The XbnAb Cohort: 305 people with broadly neutralizing antibody activity to HIV-1

**DOI:** 10.1101/2024.09.13.612733

**Authors:** Chloé Pasin, Daniel Schmidt, Merle Schanz, Baptiste Elie, Nikolas Friedrich, Irene A. Abela, Katharina Kusejko, Cyrille Niklaus, Michèle Sickmann, Jacqueline Weber, Michael Huber, Amapola Manrique, Andri Rauch, Alexandra Calmy, Matthias Cavassini, Marcel Stöckle, Julia Notter, Enos Bernasconi, Dominique L. Braun, Huldrych F. Günthard, Roger D. Kouyos, Peter Rusert, Alexandra Trkola, the Swiss HIV Cohort Study

## Abstract

Broadly neutralizing antibodies (bnAbs) recognizing a diversity of HIV-1 strains are widely thought to be essential for an HIV-1 vaccine. Extensive knowledge on bnAbs has been gained from studying natural HIV infection by following bnAb evolution in individual people with HIV (PWH). However, it remains essential to increase knowledge of bnAb responses in large PWH cohorts to assess the feasibility of inducing bnAb activity by vaccination. To allow systematic analysis, we created the XbnAb cohort, a large bnAb-inducer cohort selected by screening plasma of PWH enrolled in the Swiss HIV Cohort Study (SHCS) and the Zurich Primary HIV Infection Study (ZPHI). The XbnAb cohort represents a retrospective, biobank-based cohort comprising data of 305 PWH who developed bnAb activity during HIV-1 infection. Here, we report on the characteristics of the XbnAb cohort and its potential for HIV vaccine research.

## Introduction

A globally effective preventive vaccine remains critical to ending the HIV-1 epidemic^1,2^. Broadly neutralizing antibody (bnAb) vaccine development has entered a new era with the recent success of novel HIV-1 envelope (Env) glycoprotein-derived immunogens, including multiple-epitope, epitope-focused and germline targeting immunogens, in inducing bnAb precursors^3–16^. While a few approaches have succeeded in triggering precursors of bnAbs of different specificities, and some even reached notable neutralization capacity, sophisticated immunization regimens must still be developed to guide the evolution of these strain-specific precursors to acquire neutralizing breadth and potency needed^5,9^. The coverage and potency that an effective HIV-1 vaccine must achieve are unprecedented among established antiviral vaccines, as complete inhibition of HIV-1 transmission is needed to prevent HIV integration and seeding of a latent reservoir^1,17^. The Antibody Mediated Prevention (AMP) trial that tested passive immunization of a CD4 binding site (CD4bs) bnAb, VRC01, provided an estimate of the benchmark neutralizing activity that a preventative vaccine must achieve^18–20^. Maintaining such high levels of activity for years following vaccination will be another challenge for HIV vaccines to overcome. A combination of immune activities elicited by vaccines - either several types of bnAbs or a combination of bnAbs and cellular immune responses - are therefore recognized as critical. In combination, bnAbs may achieve the desired efficacy and coverage across HIV-1 subtypes at lower titers^21–24^ supporting the design of vaccine regimens that combine individual immunogens to induce multi-specific bnAb activity. As a result, the next challenge in bnAb vaccine development will be to determine which combined bnAb specificities a vaccine should induce, whether the different bnAb activities can be stimulated simultaneously, or whether sequential vaccine approaches will be necessary.

The polyclonal antibody response to the HIV envelope trimer, both in natural infection and after immunization, may provide critical chaperone activity for the evolution of distinct bnAbs, but may also lead to interference that limits the induction of desired responses^25–28^. Antibody lineages that target overlapping epitopes and co-evolve with bnAbs suggest a role for antibody cooperativity in the induction of bnAbs^29–32^. Equally, it may not be a single bnAb but multiple bnAb lineages with different specificities that provide the elite potency and breadth recorded as bnAb activity in plasma of some people with HIV (PWH)^33–40^. Of relevance to vaccine design, it has been shown that multiple bnAb lineages with modest breadth can provide high coverage and activity lifting the barrier of escape ^30,36,39–41^.

Increased knowledge of bnAb responses in natural infection will be essential to assess the feasibility of inducing multiple bnAb activities by vaccination. To allow a systematic analysis, we created the XbnAb cohort, a retrospective, biobank-based cohort of PWH (*n=*305) who developed bnAb activity during natural infection. Here, we report on the characteristics of the XbnAb cohort and its potential for HIV vaccine research.

## Results

We previously identified 239 PWH with bnAb activity in the Swiss 4.5K screen^42–44^ by assessing the neutralization breadth of biobanked plasma from 4,484 PWH participating in two longitudinal cohorts, the Swiss HIV Cohort Study (SHCS)^45^ and the Zurich Primary HIV Infection Study (ZPHI)^46^. The Swiss 4.5K screen leveraged a comparatively small multi-subtype HIV-1 pseudovirus panel (8-virus panel; Supplementary Table 1) to rank plasma neutralization activity into categories of no/low, cross, broad and elite neutralization activity, with the latter two categories comprising the 239 participants denoted as bnAb inducers. Here, 498 plasma from the Swiss 4.5K screen that showed cross-neutralizing activity against the 8-virus panel but did not meet the bnAb inducer threshold were re-assayed with a larger, 23-virus panel, identifying 65 additional bnAb inducers (Supplementary Figure 1 and methods). A further bnAb inducer was identified by screening against a 41-virus panel accounting for a total of 305 bnAb inducers that were assembled into the XbnAb plasma cohort (Figure 1). The XbnAb cohort includes bnAb inducers with moderate to high neutralization breadth (median 61%, 1Q-3Q=49%-73%) and potency (median geometric mean of ID50=420, 1Q-3Q=340-567) as defined against a 41 multi-subtype virus panel (Supplementary Table 1 and 2). Demographic and disease parameters in the XbnAb cohort mirror those reported for bnAb inducers in the Swiss 4.5K screen^44^ with parameters that reflect exposure to HIV-1 antigen (length of untreated infection, viral load and diversity) and black ethnicity being independently associated with bnAb evolution compared to PWH without bnAb activity (Figure 2, Table 1, Supplementary Figures 2 and 3).

**Figure 1.**
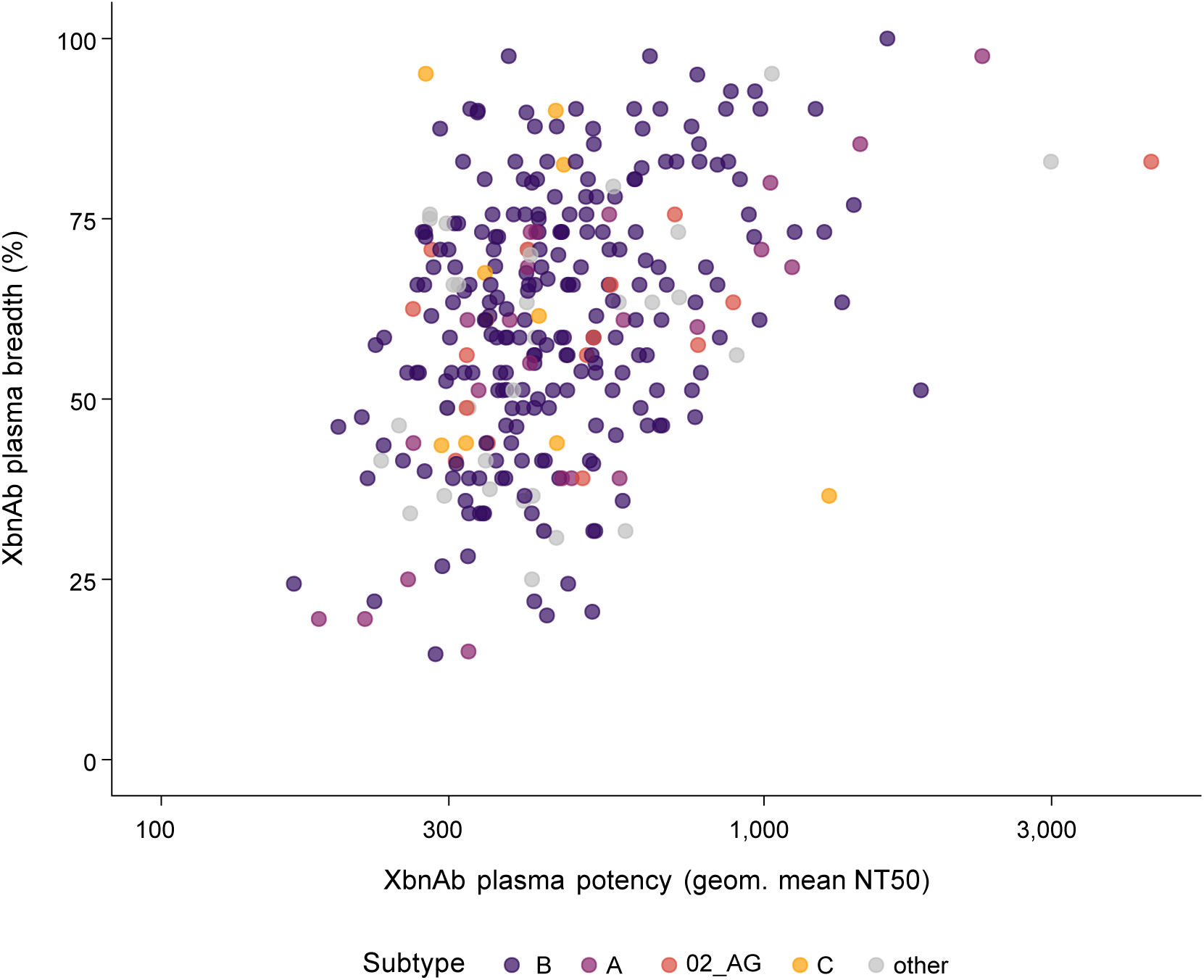
Breadth and potency in the XbnAb cohort. Plasma neutralization breadth (percentage of viruses neutralized in the 41-virus panel) versus potency (geometric mean 50% neutralization titer (NT50)) of all bnAb inducers included in the XbnAb cohort (*n*=305). Colors indicate HIV-1 *pol* subtypes of donors.

**Figure 2.**
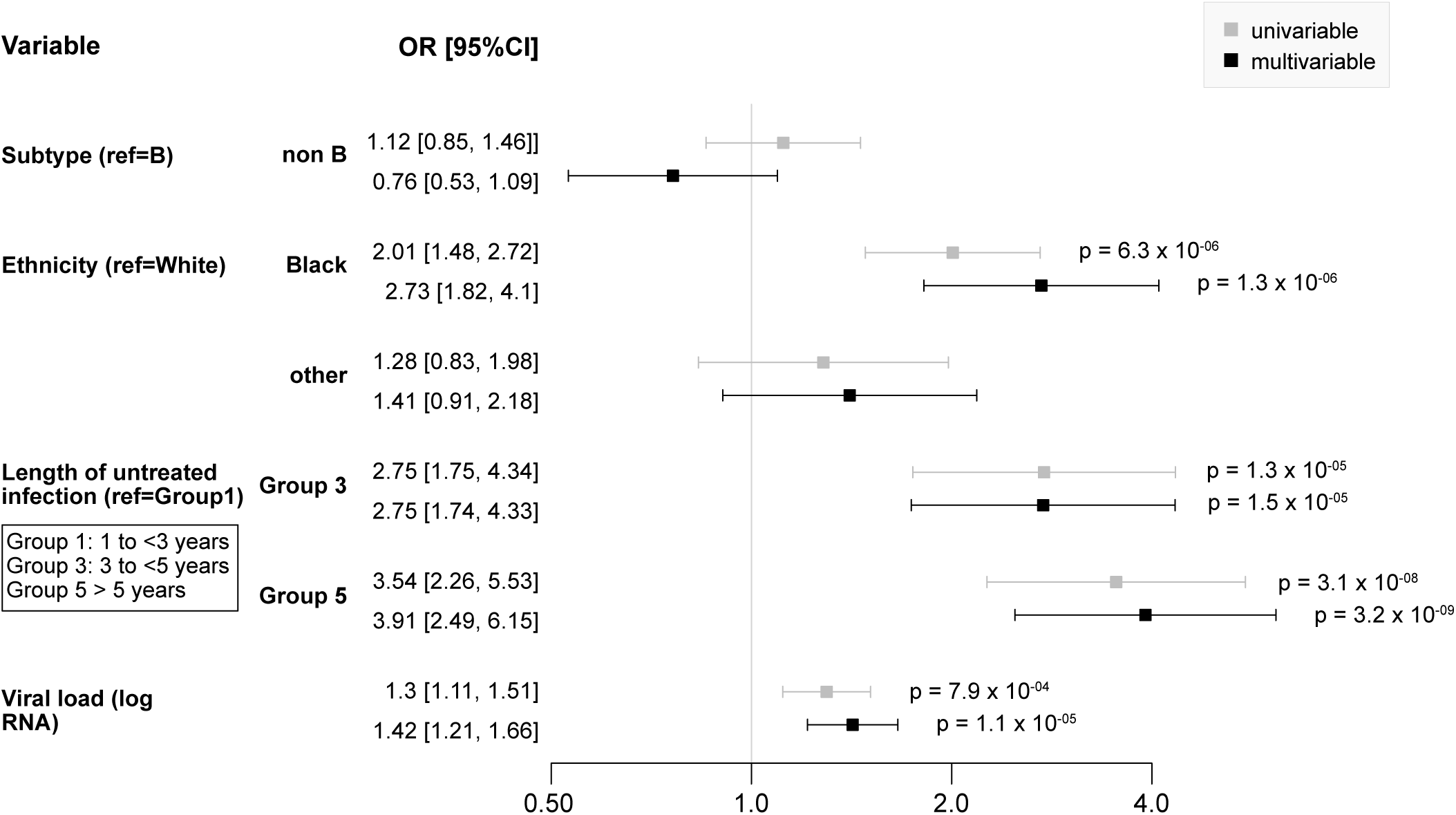
Parameters linked with bnAb status in the XbnAb cohort. Association between demographic and disease parameters with bnAb inducer status, as estimated with odds ratios (OR) in a logistic regression comparing bnAb inducers in the XbnAb cohort (n=305, identified through the Swiss 4.5K screen) with the remaining participants of the Swiss 4.5K screen not classified as bnAb inducers (n=4179). Analyzed variables derived from⁴⁴ included: HIV-1 pol subtype, ethnicity (race), length of untreated HIV infection (Group 1: infected >1<3 years, Group 3: infected 3 > 5 years, Group 3: infected > 5 years). Participants with available data for all variables were included; XbnAb cohort (n=294/305), Swiss 4.5K screen participants without bnAb activity (n=3692/4179).

**Figure 3.**
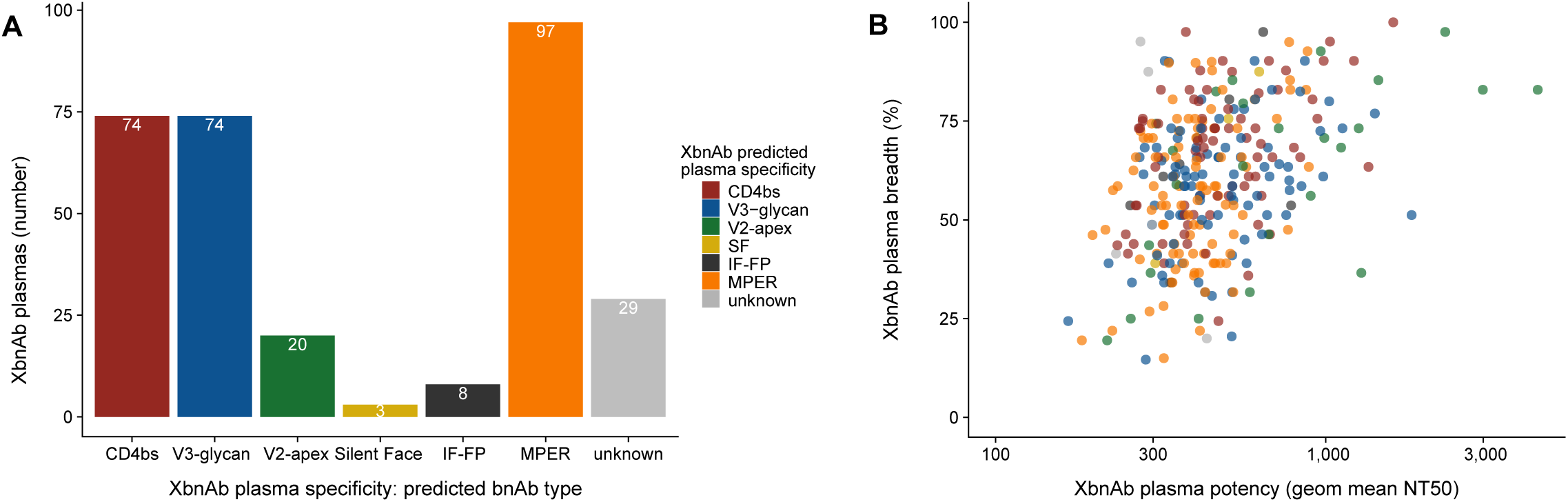
Specificity, breadth and potency of bnAb activity in the XbnAb cohort. **A.** Plasma specificities of the XbnAb donors as determined by neutralization specificity prediction through maximum Spearman based prediction (MSBP) using plasma neutralization data against the 41-virus panel. Colors indicate predicted plasma specificities. **B.** Plasma neutralization breadth (percentage of viruses neutralized in the 41-virus panel) versus potency (geometric mean ID50) of all bnAb inducers in the XbnAb cohort (n=305). Colors indicate predicted bnAb type.

**Table 1:**
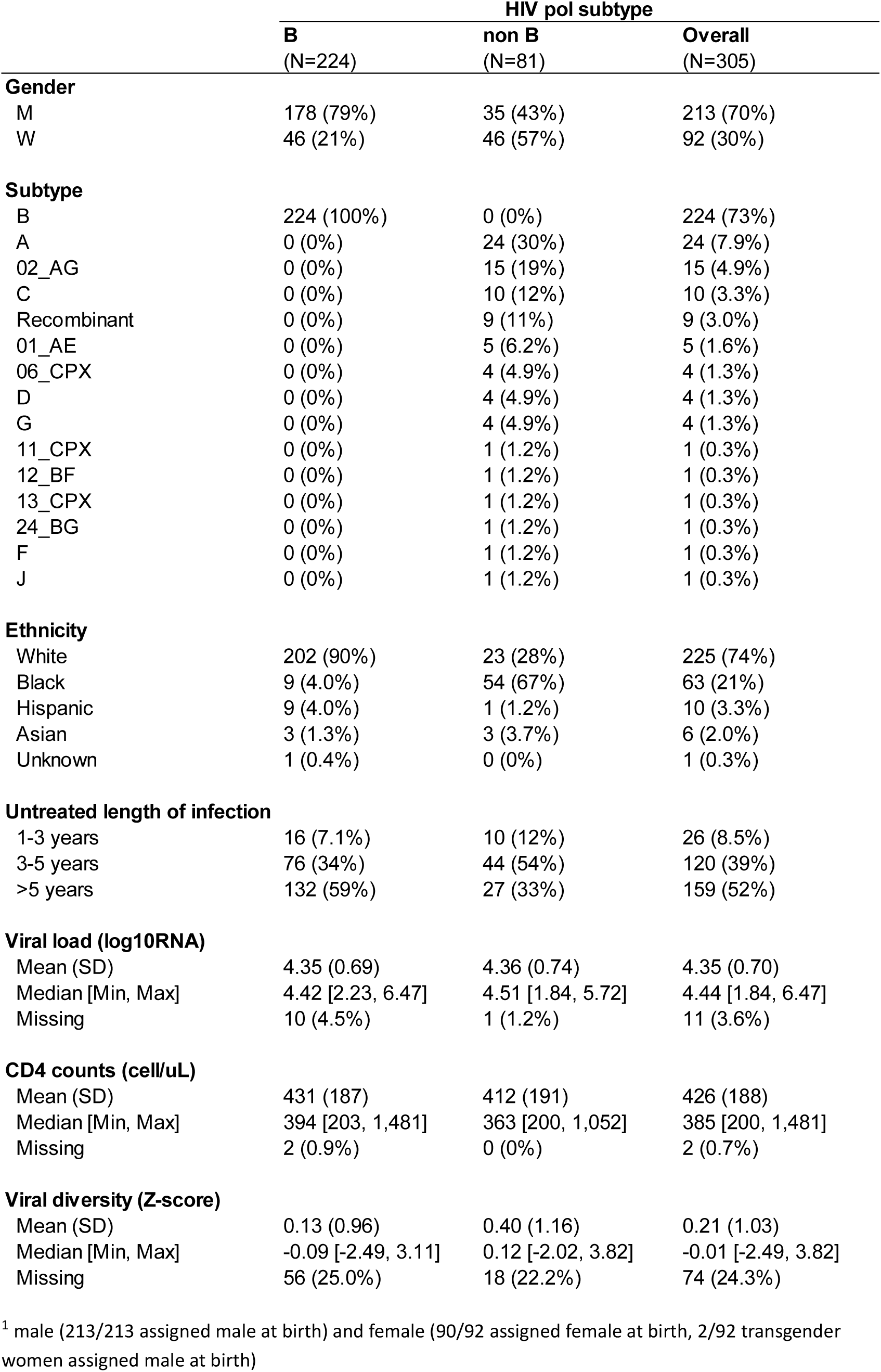
Demographic and disease parameters in the XbnAb cohort.

Definition of the bnAb specificity in the XbnAb cohort based on comparison of neutralization fingerprints on the 41-virus panel with reference bnAbs (*n=*46, Supplementary Table 3) using a Spearman-correlation based approach^44^ was successful in predicting a dominant bnAb epitope specificity for 276/305 of the bnAb inducers (Figure 3). We recorded bnAb activity of all main bnAb categories identified thus far (CD4 binding site (CD4bs), V2-apex, V3-glycan, silent face (SF), interface/fusion peptide (IF-FP), membrane proximal external region (MPER)). Consistent with our previous observations, analysis of the XbnAb cohort confirmed a subtype preference of CD4bs and V2 bnAb activity, with CD4bs activity more common in subtype B (Fisher test, p=0.0098) and V2 bnAb activity more common in non-B infection (Fisher test, p=3.2×10^−5^; Figure 4).

**Figure 4.**
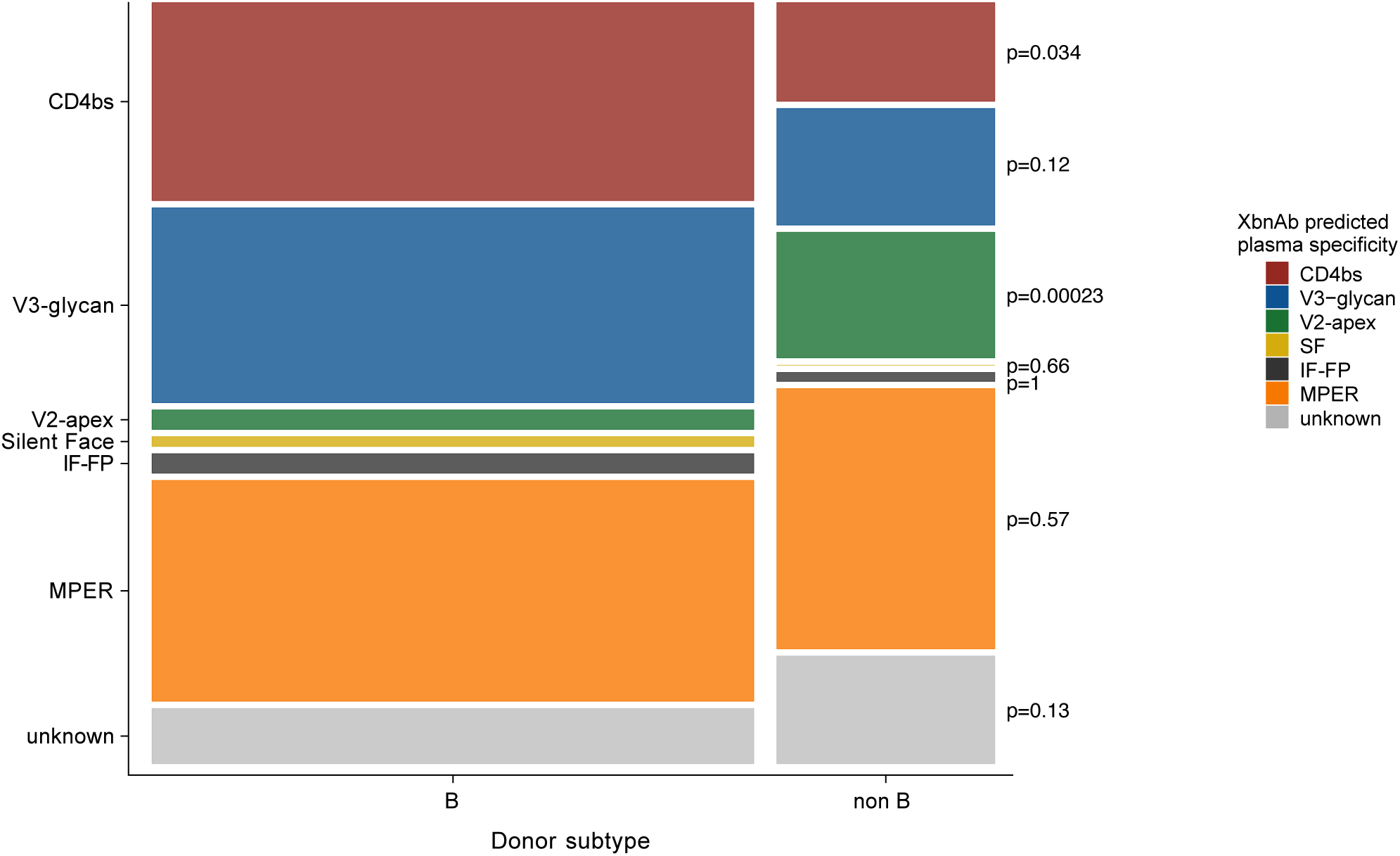
Association between HIV-1 subtype and bnAb specificity in the XbnAb cohort. HIV-1 subtype was defined through available information in the SHCS and ZPHI databases on HIV-1 *pol* subtype and classified as subtype B versus non-B. Association between subtype and plasma bnAb specificity was estimated by Fisher’s exact test. Rectangle sizes correspond to the proportion of plasma samples in each sub-category.

We conclude that the expanded number of bnAb inducers in the XbnAb cohort fully represents the patterns observed in the original Swiss 4.5K screen, allowing its use as a resource for in-depth investigation of the parameters of bnAb evolution.

## Discussion

A key strength of the XbnAb cohort is the breadth and longitudinal depth of the data accrued over decades of follow-up, together with its derivation from a large underlying population of nearly 4500 PWH. An important resource leveraged in the Swiss 4.5K study, and further developed within the XbnAb cohort, is the extensive dataset of viral, demographic, and clinical parameters collected through the SHCS and the ZPHI.

HIV-1 infections in Switzerland are dominated by Subtype B with sizeable proportion of diverse non-B subtypes^47–50^. The Swiss HIV Cohort Study (SHCS) includes 71% of people with HIV who are on antiretroviral therapy (ART) in Switzerland. The study population is predominantly white (67.8%), male (72.7%), and men who have sex with men (MSM) (39.2%) ^45^. The unique composition of the SHCS, with this primary population and a significant proportion of PWH from diverse demographics, offers an opportunity to assess parameters that influence HIV-1 disease with high statistical power in diverse settings, including different genders, different transmission risk groups, and different HIV subtypes^47,51–54^. This capacity extends to the XbnAb cohort, as we show here.

Both the SHCS and the ZPHI study cohorts provide opportunities for recruiting participants into clinical trials that can be also leveraged for XbnAb donors that are still actively enrolled in these cohorts. A first vaccination study, RENEW-SHCS (SNCTP000006192), recruiting XbnAb participants is currently conducted^55^. This study builds on the recently introduced concepts of immunizing PWH with bnAb immunogens to evaluate their efficacy in stimulating bnAb activity and to guide further immunogen development^2,56^.

## Methods

### Human Specimen and Ethics

The XbnAb cohort, as referred to here, is a biobank cohort, consisting of a collection of plasma samples from PWH who were defined to harbor bnAb activity in the Swiss 4.5K screen^42–44^. The XbnAb cohort constitutes of plasma samples provided for the Swiss 4.5K screen by two active recruiting cohorts, the Swiss HIV Cohort study (SHCS)^45,57^ and the Zurich Primary HIV Infection Study (ZPHI)^46^, which maintain extensive data and biobanks. We refer to the XbnAb cohort as cohort, as the analyzed plasma samples are connected to active cohorts, the SHCS and the ZPHI, and a sizeable proportion of the participants is still actively enrolled in their respective recruiting cohorts.

The SHCS is registered under the Swiss National Science longitudinal platform program https://data.snf.ch/grants/grant/201369). The SHCS is a prospective, nationwide, longitudinal, observation, clinic-based cohort enrolling PWH in Switzerland since 1988. Participants are followed-up biannually with clinical visits and blood collection. The ZPHI is an ongoing, observational, nonrandomized, single center cohort founded in 2002 that specifically enrolls patients with documented acute or recent primary HIV-1 infection (ClinicalTrials.gov identifier NCT00537966).

The SHCS and the ZPHI studies were approved by the ethics committees of the participating institutions (Kantonale Ethikkommission Bern, Ethikkommission des Kantons St. Gallen, Comite Departemental d’Ethique des Specialites Medicales et de Medicine Communataire et de Premier Recours, Kantonale Ethikkommission Zürich, Repubblica et Cantone Ticino–Comitato Ethico Cantonale, Commission Cantonale d’Étique de la Recherche sur l’Être Humain, Ethikkommission beider Basel for the SHCS and Kantonale Ethikkommission Zürich for the ZPHI), and written informed consent was obtained from all participants.

### XbnAb cohort

The XbnAb cohort is based on a retrospective analysis of biobanked plasma. It is a collection of plasmas of 305 bnAb inducers identified through the Swiss 4.5K screen^42–44^. Participants were not actively recruited into the XbnAb cohort. They entered their recruiting cohorts at different stages of their infection and were followed there longitudinally as long as participants remained enrolled. The Swiss 4.5K screen^42–44^ leveraged biobanked plasma from antiretroviral-free periods of participants reaching back to the foundation of the SHCS in 1988 and the ZPHI in 2002. The Swiss 4.5K screen specifically included plasma samples collected during periods without ART and stratified participants according to length of untreated infection into groups with more than 1, more than 3, and more than 5 years of untreated HIV infection. This plasma screening timepoint remained the point of reference for the XbnAb cohort and is used to classify neutralization activity and bnAb inducer status. The 305 bnAb inducers that form the XbnAb cohort comprise 239 bnAb inducers identified in the original Swiss 4.5K screen by reaching a plasma neutralization score > 9 against an 8-virus panel as described in Rusert et al. ^44^ (Supplementary Table 1). To identify additional bnAb inducers, plasmas from 499 donors that in the first screen either reached a score of 7-9, or neutralized at least one virus at >80%, were re-screened on a 23-virus panel (Supplementary Table 1). BnAb inducer plasmas that reached a score of >26 in the 23-virus screen (*n=*65) were included into the XbnAb cohort. One additional donor was identified as bnAb inducer by direct screening against a 41-virus panel accounting for the final size of 305 bnAb inducers. The fully assembled 305 bnAb inducer plasma collection was subjected to plasma fingerprinting using a 41-virus panel and a reference bnAb panel as defined below. In addition to the plasma neutralization data the XbnAb cohort utilizes demographic, host and viral data collected by the SHCS and ZPHI as described in the Swiss 4.5K screen^42–44^.

### Cell lines

HEK 293T cells (American Type Culture Collection, USA) and TZM-bl cells (NIH AIDS Reagent Program, USA) were cultivated in DMEM, high glucose, pyruvate supplemented with 10% heat-inactivated FBS, 100 U ml−1 penicillin and 100 µg ml−1 streptomycin (all from Gibco, Thermo Fisher Scientific, USA) at 37 °C, 5% CO2 and 80% relative humidity.

### HIV-1 Pseudovirus panels

Plasma neutralization activity in the XbnAb cohort was assessed using multi-subtype HIV-1 Envelope (Env) pseudovirus panels. A full list of viruses included in each panel is available in their Env sources is listed in Supplementary Table 1. The initial neutralization screen was conducted in the Swiss-4.5K screen using an 8-virus panel^44^. Re-screening of 499 cross-neutralizing plasmas to identify additional bnAb inducers was done using a 23-virus panel. One Swiss 4.5K donor was identified as bnAb inducer by direct screening against a 41-virus panel. The 41-virus panel was used to define the final neutralization breadth and potency of the XbnAb cohort (*n=*305) and for neutralization fingerprinting. The 41-virus panel contains Env pseudoviruses of diverse subtypes and circulating recombinant forms (subtype B, *n=*14; subtype C, n=11; subtype A, *n=* 6; recombinant form AE, *n=*5; subtype G, *n=*3; recombinant forms ACD and AG, *n=*1).

### Reference bnAbs

We used a set of 46 bnAbs with defined target epitopes as detailed in Supplementary Table 3 as a reference for comparison of neutralization fingerprints with PWH plasma. 50% inhibitory concentrations (IC50, µg/ml) of the reference bnAbs against the 41-virus panel are listed in Supplementary Table 3.

### HIV-1 pseudovirus neutralization

Neutralization activity was measured in a 384-well format using the Env pseudovirus luciferase reporter assay on TZM-bl cells as described^44^. Luciferase reporter activity (relative light units (RLU)) was measured on an EnVision Multilabel Reader (PerkinElmer, Inc., USA) three days post-infection using Luciferase assay reagent (Promega, USA) as described^44^. Plasma was heat-inactivated before use in the neutralization assay. Plasma and bnAb dilutions reported refer to the final assay volume including cells.

The <u>23-virus neutralization screen</u> was conducted analogous to the 8-virus screen described in Rusert et al. ^44^. A 1/150 dilution of heat-inactivated plasma was preincubated with the respective virus for 1 h before infection of target cells. Neutralization activity was calculated as the reduction in infectivity measured as RLU per well compared to infection in absence of plasma or bnAb. Plasma neutralization activity was scored as follows: score=0 for neutralization activity below 20%, score=1 for neutralization activity between 20% and <50%, score=2 for neutralization between 50% and <80%, score=3 for neutralization ≥80%. Plasmas were then ranked by the sum of scores against all 23 viruses to reflect their potency and breadth. The maximum cumulative neutralization score that could be achieved by a plasma was 69. Plasmas with a score >26 were included into the XbnAb cohort.

<u>41-virus screen</u>: Serial dilutions of bnAbs or plasma (starting at dilution 1/100) were measured to define the plasma or antibody concentrations causing 50% reduction in viral infectivity (half-maximum inhibitory concentration, IC50 or half-maximum neutralization titer, NT50). IC50/NT50 values were calculated by fitting data to sigmoid dose–response curves (variable slope) using Prism (GraphPad Software, Inc., USA). If 50% inhibition was not achieved at the highest or lowest inhibitor concentration, a greater than or less than value was recorded. IC50 and NT50 concentrations correspond to the final assay volume.

Plasma breadth was computed as the number of neutralized (NT50>100) viruses on the 41-virus reference panel. Potency was computed as the geometric mean of neutralization titers on the 41-vvirus panel.

### Spearman-based neutralization fingerprinting

Plasma neutralization fingerprints of bnAb inducers were evaluated using the 41-virus multiclade panel and compared to the reference bnAbs, using a Spearman correlation approach (maximum spearman based prediction, MSBP) that records the maximal correlation with a specific bnAb type as described in^44^. With this fingerprinting method, the neutralization specificity of a plasma was assessed by identifying the bnAb with the most similar neutralization profile. Similarity was measured using the Spearman correlation of the IC50 values of the bnAbs and 1/NT50 values of the plasmas. Plasmas with Spearman correlation lower than 0.4 with all reference bnAbs were classified as “unknown”.

### Statistical analyses

Statistical analyses were conducted in R version 4.4.1. To compare the XbnAb cohort to the rest of the Swiss 4.5K screen, we used logistic regressions adjusted on confounders, including subtype, ethnicity/race, evolution time (i.e., time off ART), viral load, CD4 levels, viral diversity as in Ruster et al.^44^. A Wald test with two-sided hypothesis was used to assess the significance of the estimated odds-ratio. Fisher’s exact test was used to compare the proportions of plasmas with given predicted specificities in B and non-B subtypes.

### Figure preparation

All figures were prepared using R version 4.4.1 (package ggplot2). Figures were assembled and finalized in Affinity Designer (Serif Europe Ltd, United Kingdom).

### Writing and Editing

During the preparation of this work, the authors used DeepL Translate, DeepL Write, ChatGPT 3.5 and Copilot for language editing and scite_ for literature search. After using these tools, the authors reviewed and edited the content as needed and take full responsibility for the content of the publication.

## Acknowledgements

This was work was funded by Swiss National Science Foundation (SNSF) grant 3147308_201266 (to AT) and the small nested SHCS project 744 and 745 (to AT). This study was co-financed within the framework of the Swiss HIV Cohort Study, supported by the SNF (#148522 and 201369 to HFG) and the SHCS research foundation. IAA was supported by a research grant of the Promedica Foundation.

We particularly thank the participants and the clinical and administrative staff of the ZPHI and SHCS cohorts for their decades of dedication, without which this research would not have been possible.

## Author contributions

Conceptualization: AT, HFG, RDK. Methodology: CP, PR, BE. Validation. CP, BE, PR, DS, MS, RDK. Formal analysis: CP, BE, PR, DS, MS, NF, IAA, CN, MS, JW. Investigation : CP, BE, PR, DS, MS, NF, IAA, CN, MS, JW. Resources: KK, MH, IAA, AC, MC, AR, MS, JN, EB, DL, HFG, the Swiss HIV Cohort Study. Data Curation : CP, BE, RDK, KK, PR, MS. Writing - Original Draft: AT, CP, MS, DS, RP. Writing - Review & Editing: AT, CP, BE, MS, DS, RP, HFG, RDK, AM. Visualization: CP, BE, MS, PR, DS. Supervision: AT, HFG, RDK. Project administration: AM, MS. Funding acquisition: AT, HFG

---

**Supplementary Figure 1:**
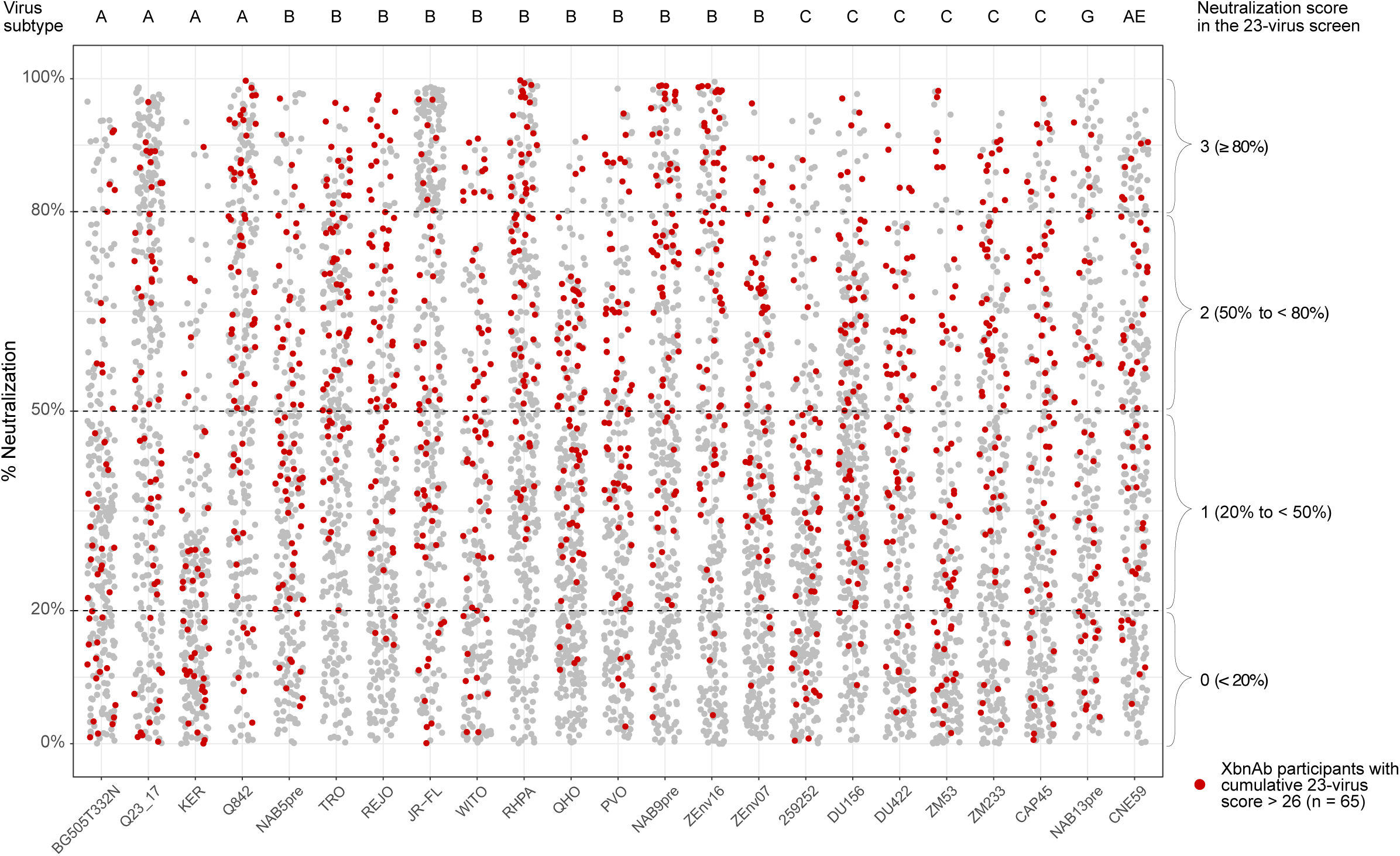
Re-screening of Swiss 4.5K donors with cross-neutralization activity with the 23-virus panel. Plasma of Swiss 4.5K donors (n=498) with neutralization scores below the threshold for bnAb inducers in the 8-virus screen^44^ but notable cross-neutralization activity (score 7-9 against 8-virus panel, or one virus neutralized > 80%) were re-screened against the 23-virus panel (plasma dilution 1/150). Percent neutralization by each plasma/virus combination was recorded and scored as indicated. Brackets indicate scoring categories of plasma neutralization activity. The maximum cumulative neutralization score against all 23 viruses that could be achieved by a plasma was 69. Plasmas with a score >26 (red) were included in the XbnAb cohort.

**Supplementary Figure 2.**
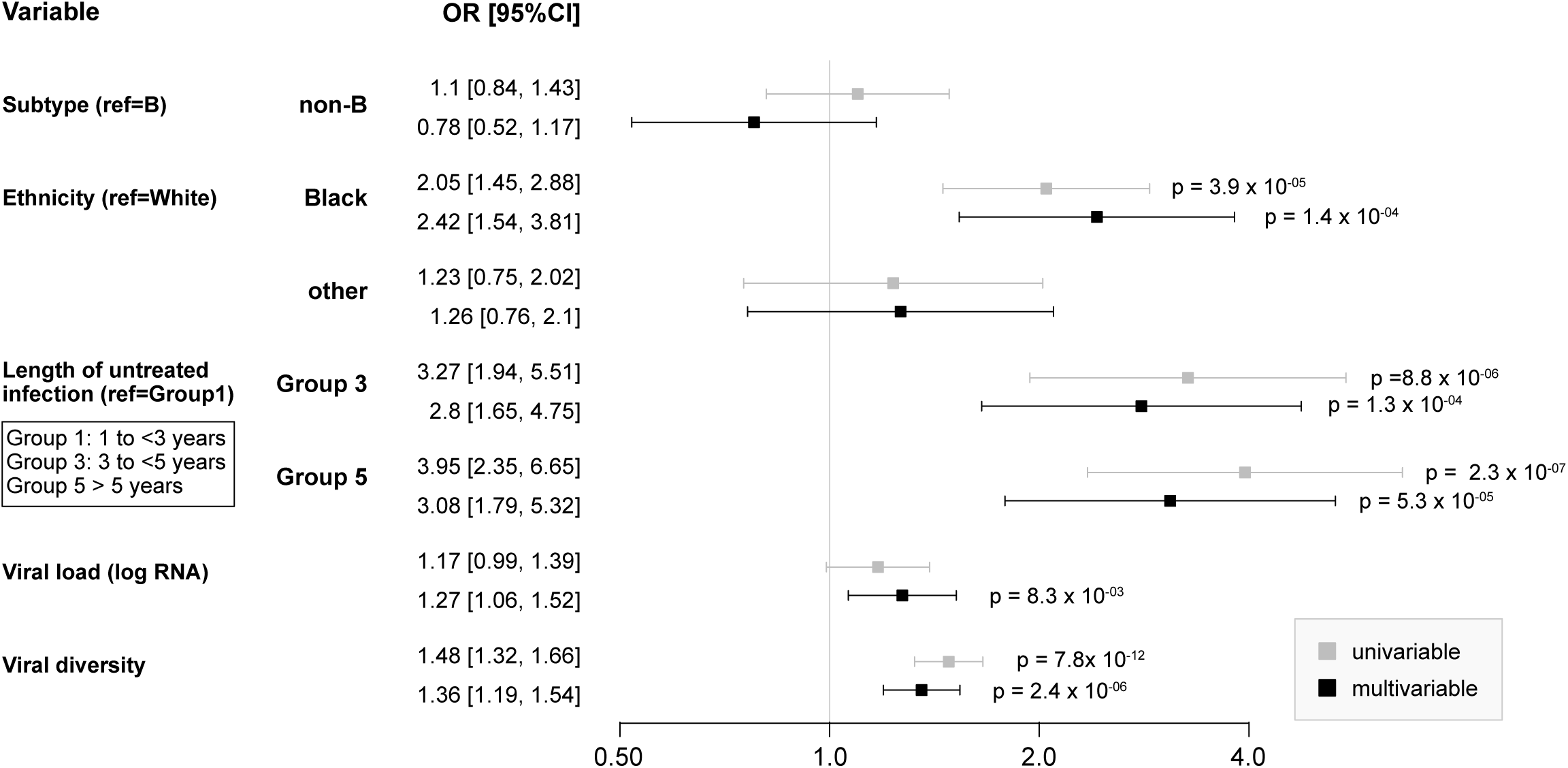
Parameters linked with bnAb status in the XbnAb cohort, including viral diversity. Association between demographic and disease parameters with bnAb inducer status, as estimated with odds ratios (OR) in a logistic regression comparing bnAb inducers in the XbnAb cohort (n=305, identified through the Swiss 4.5K screen) with the remaining participants of the Swiss 4.5K screen not classified as bnAb inducers (n=4179). Analyzed variables derived from⁴⁴ included: HIV-1 pol subtype, ethnicity (race), length of untreated HIV infection (Group 1: infected >1<3 years, Group 3: infected 3 > 5 years, Group 3: infected > 5 years), and viral diversity. Participants with available data for all variables were included; XbnAb cohort (n=224/305), Swiss 4.5K screen participants without bnAb activity (n=3040/4179).

**Supplementary Figure 3.**
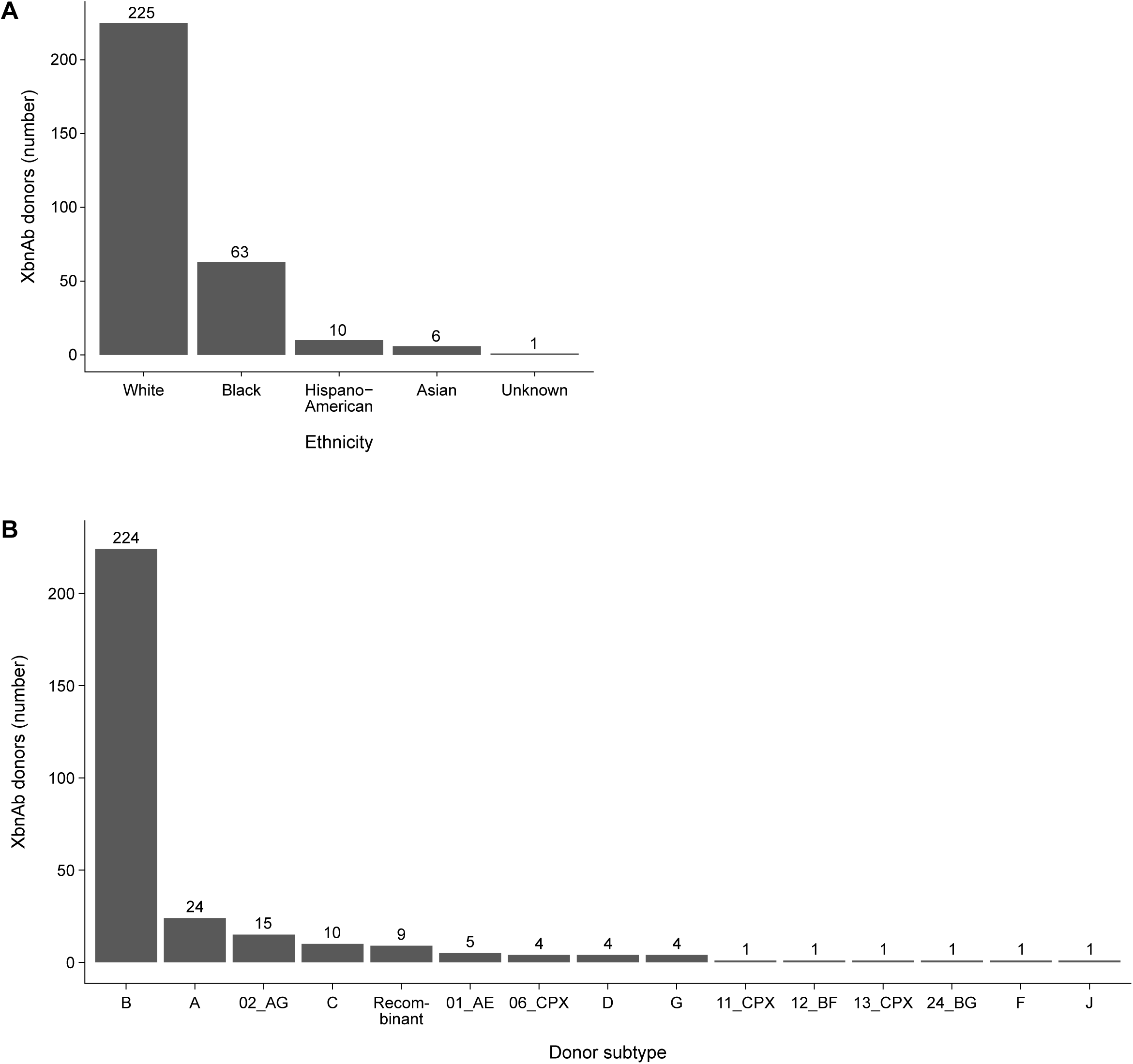
Distribution of ethnicity and subtype in the XbnAb cohort. **A.** Ethnicity of the XbnAb cohort participants as reported in the SHCS. **B.** HIV-1 *pol* subtype of the XbnAb cohort participants.

**Supplementary Table 1.**
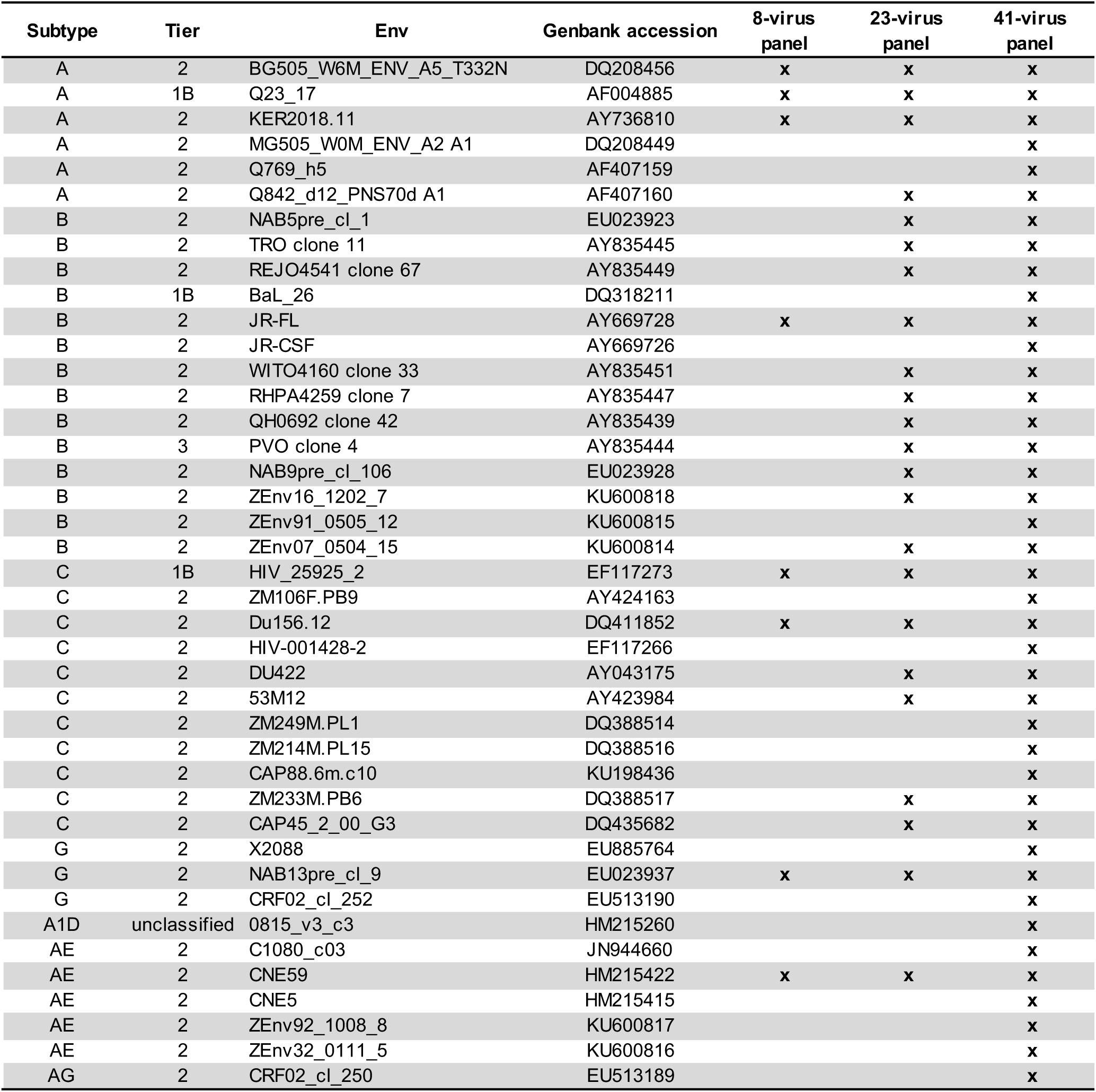
Multi-clade virus panels.

**Supplementary Table 2.**
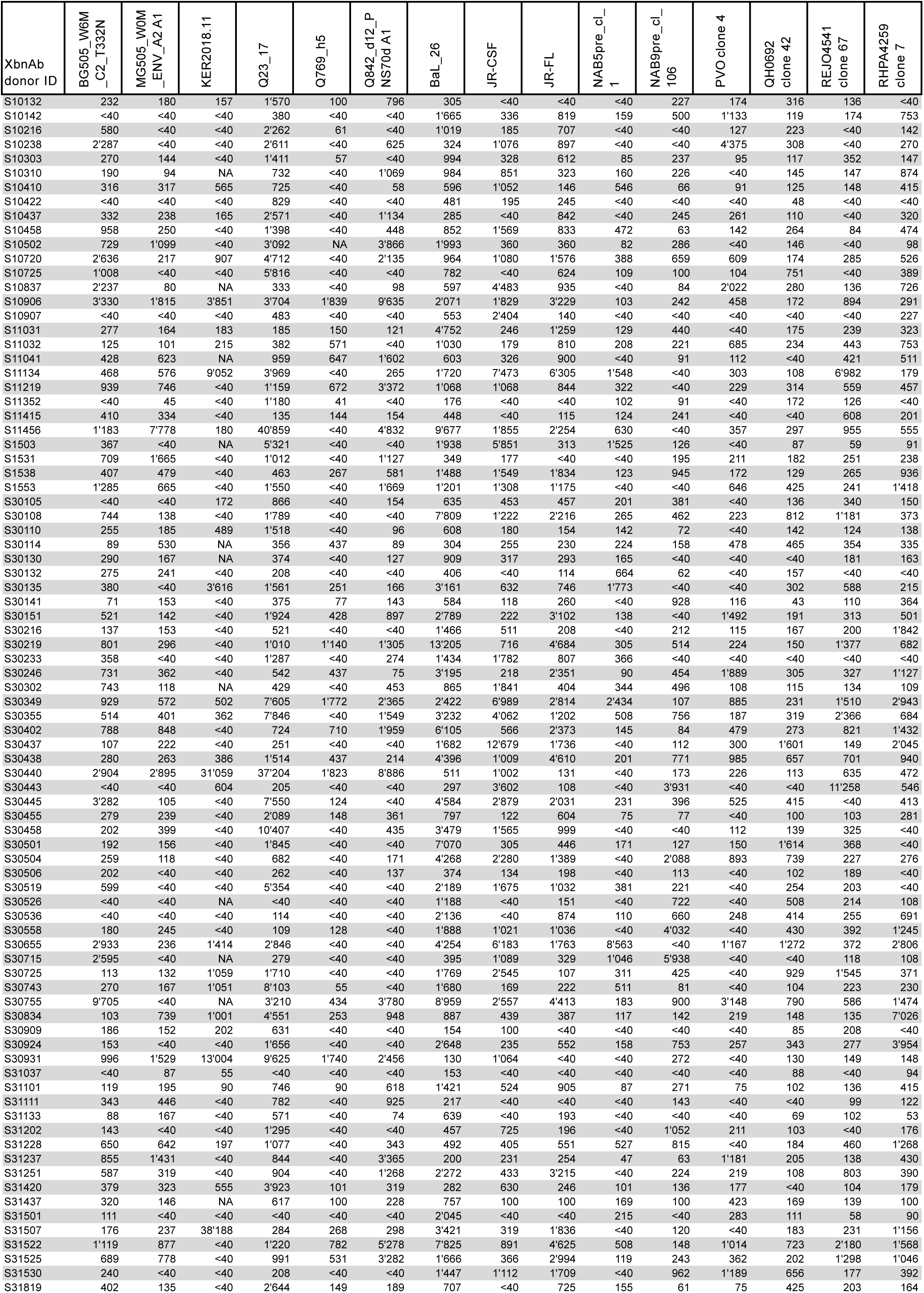
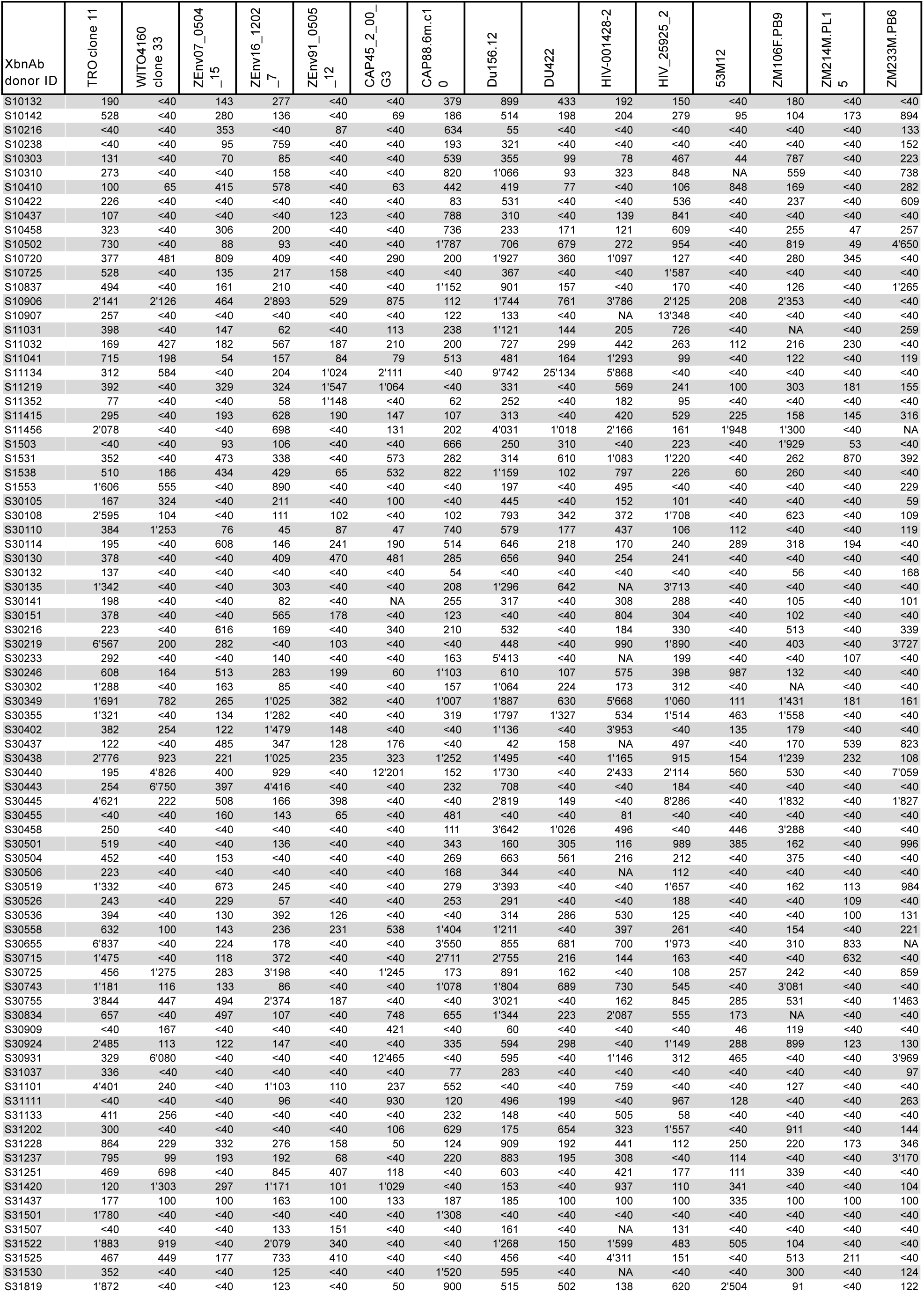
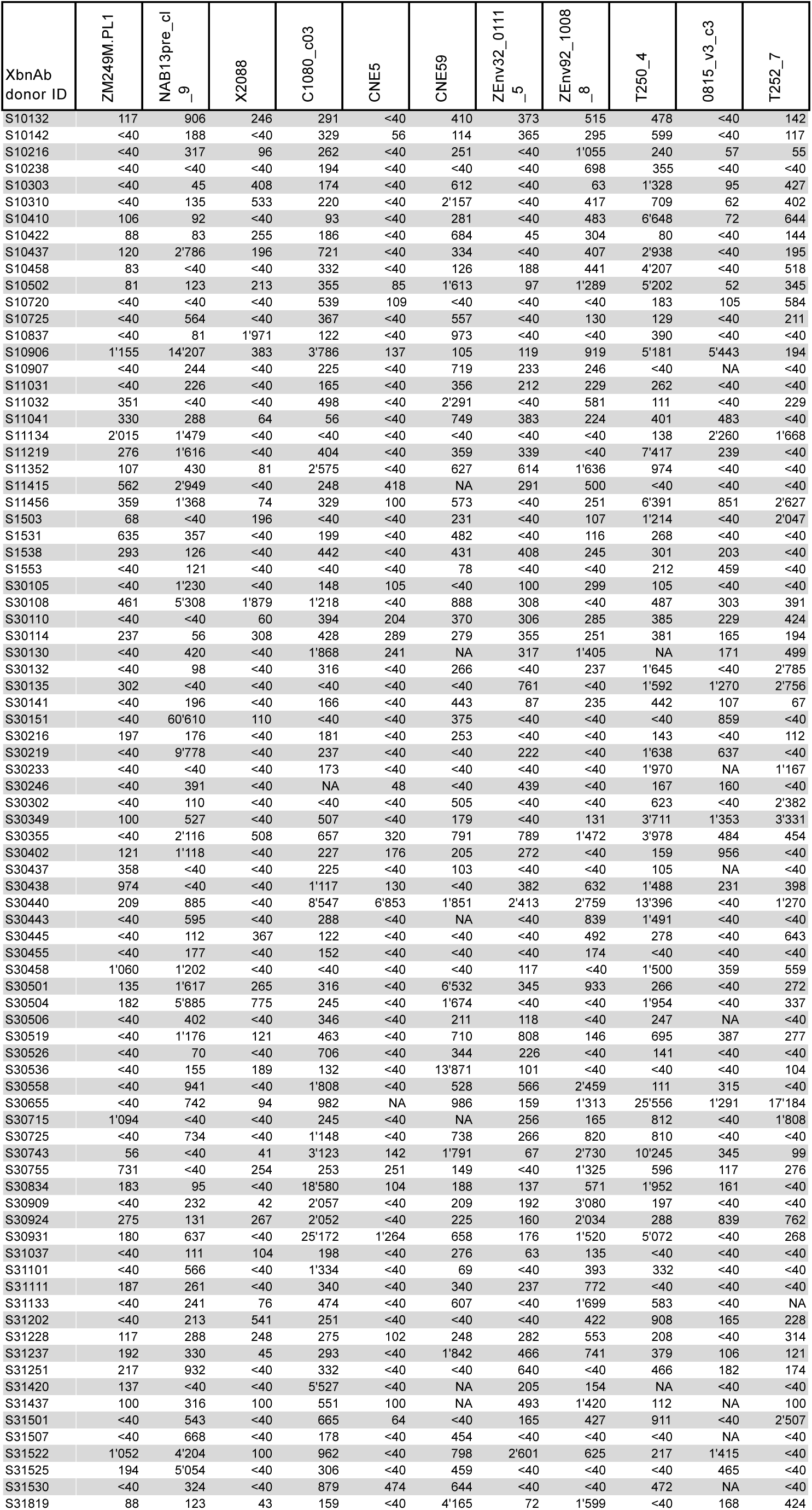
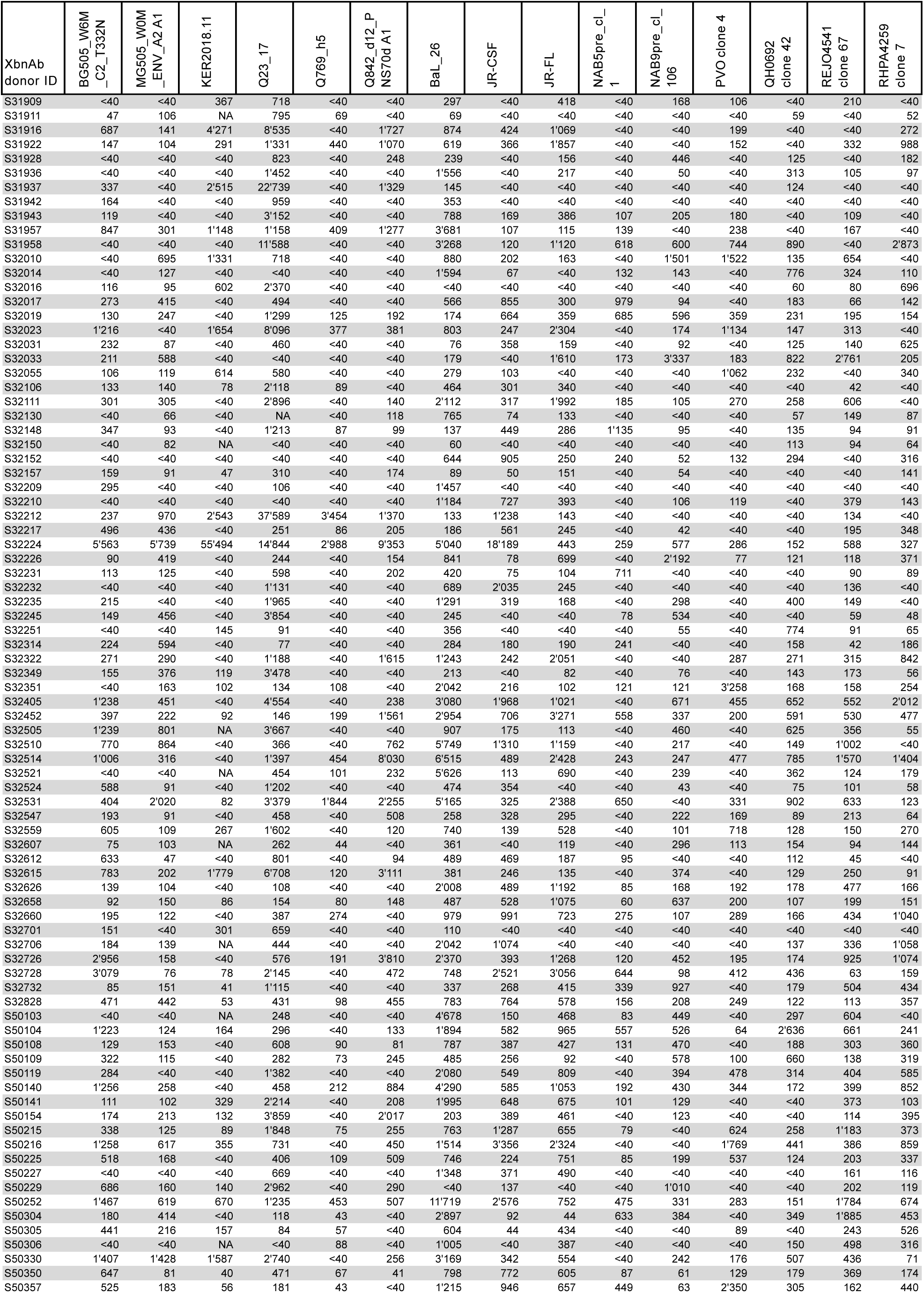
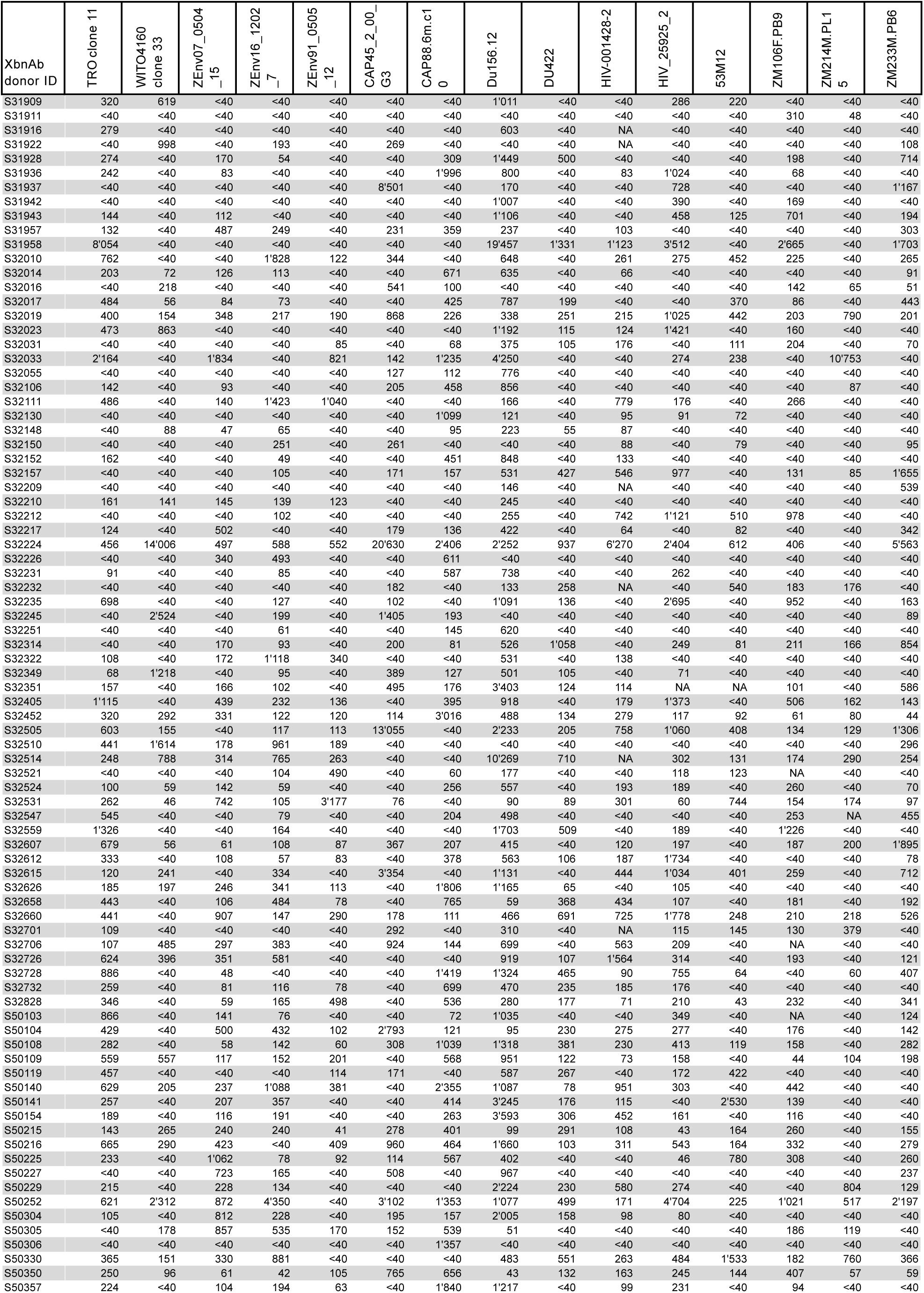
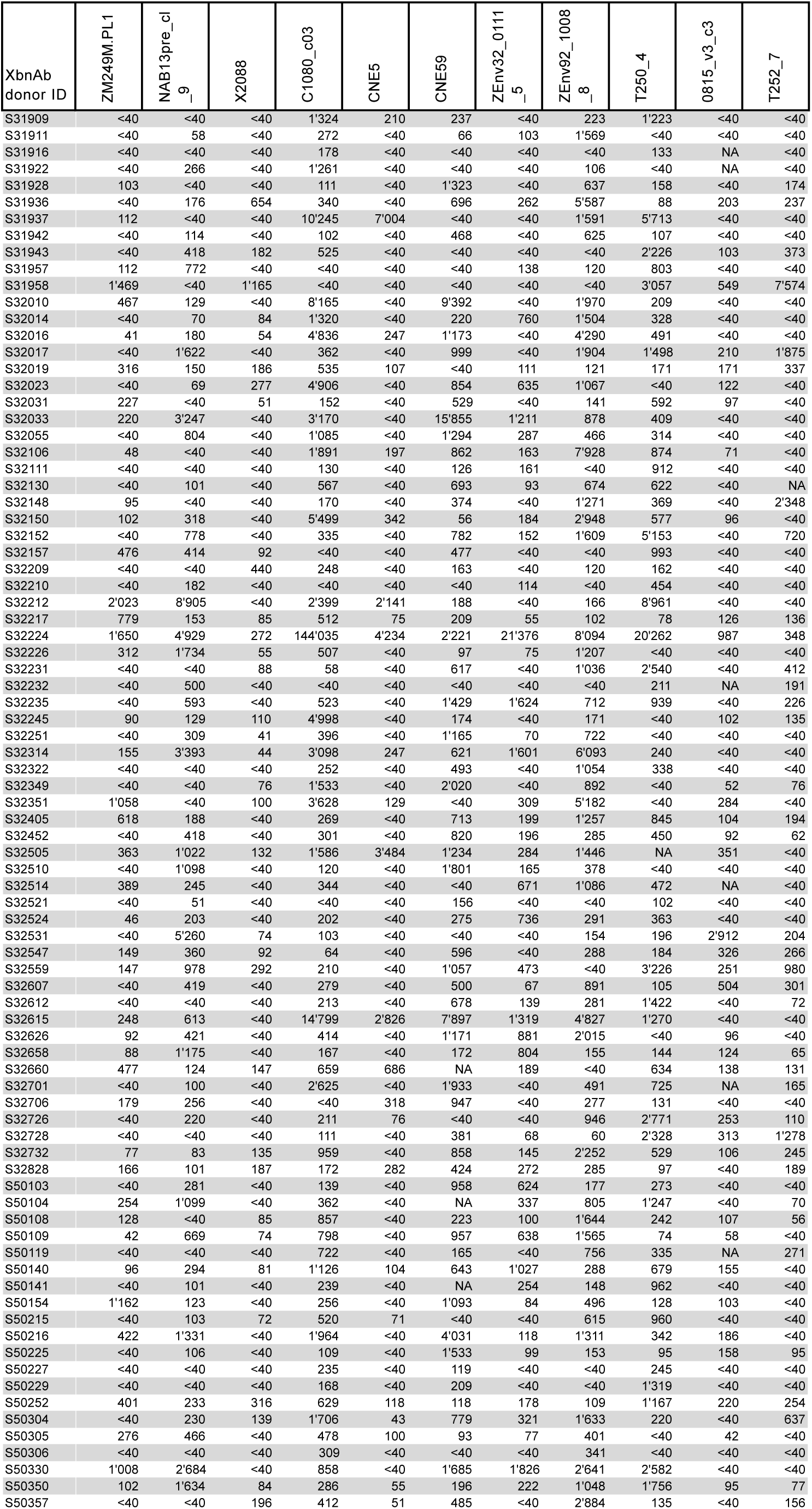
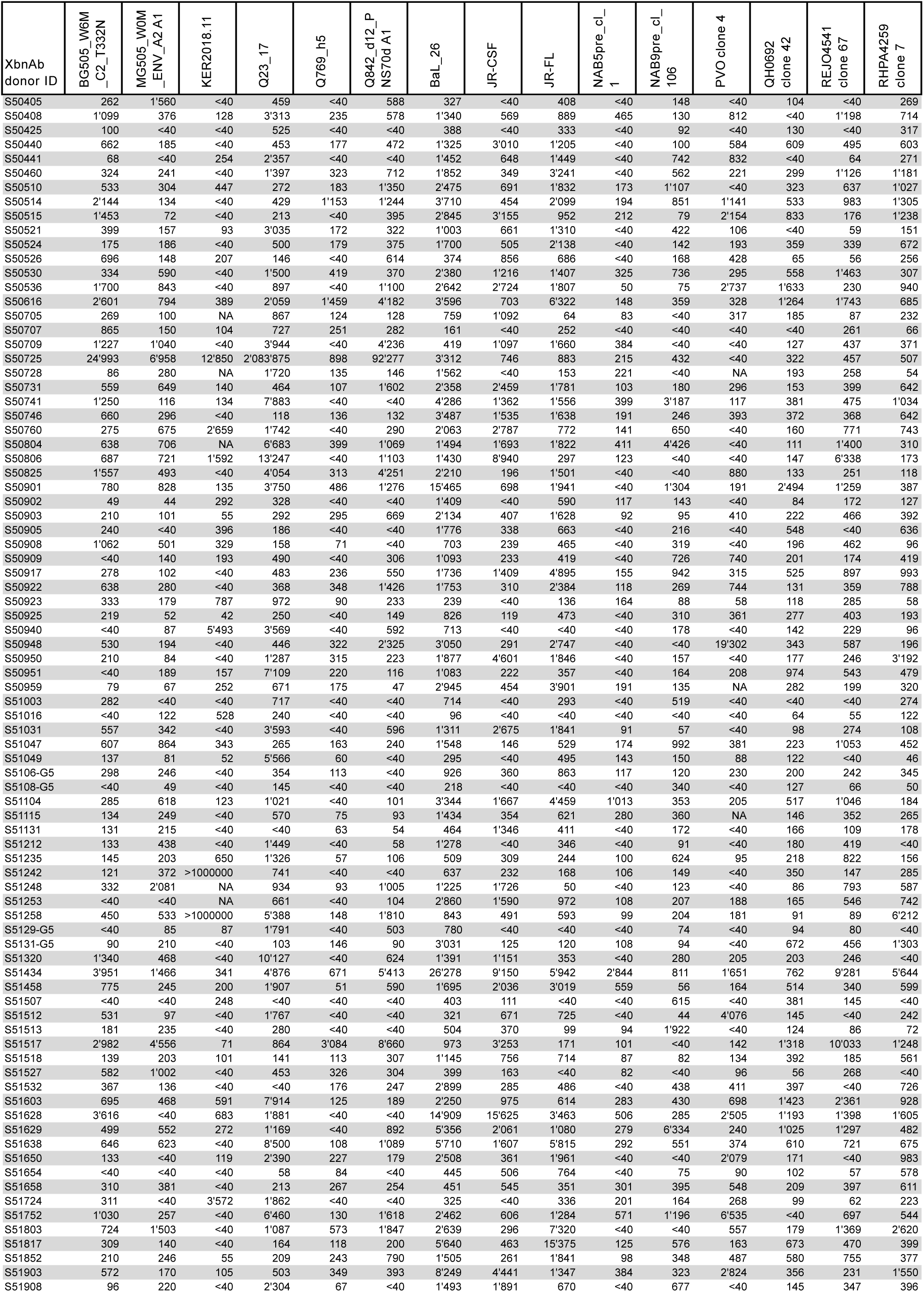
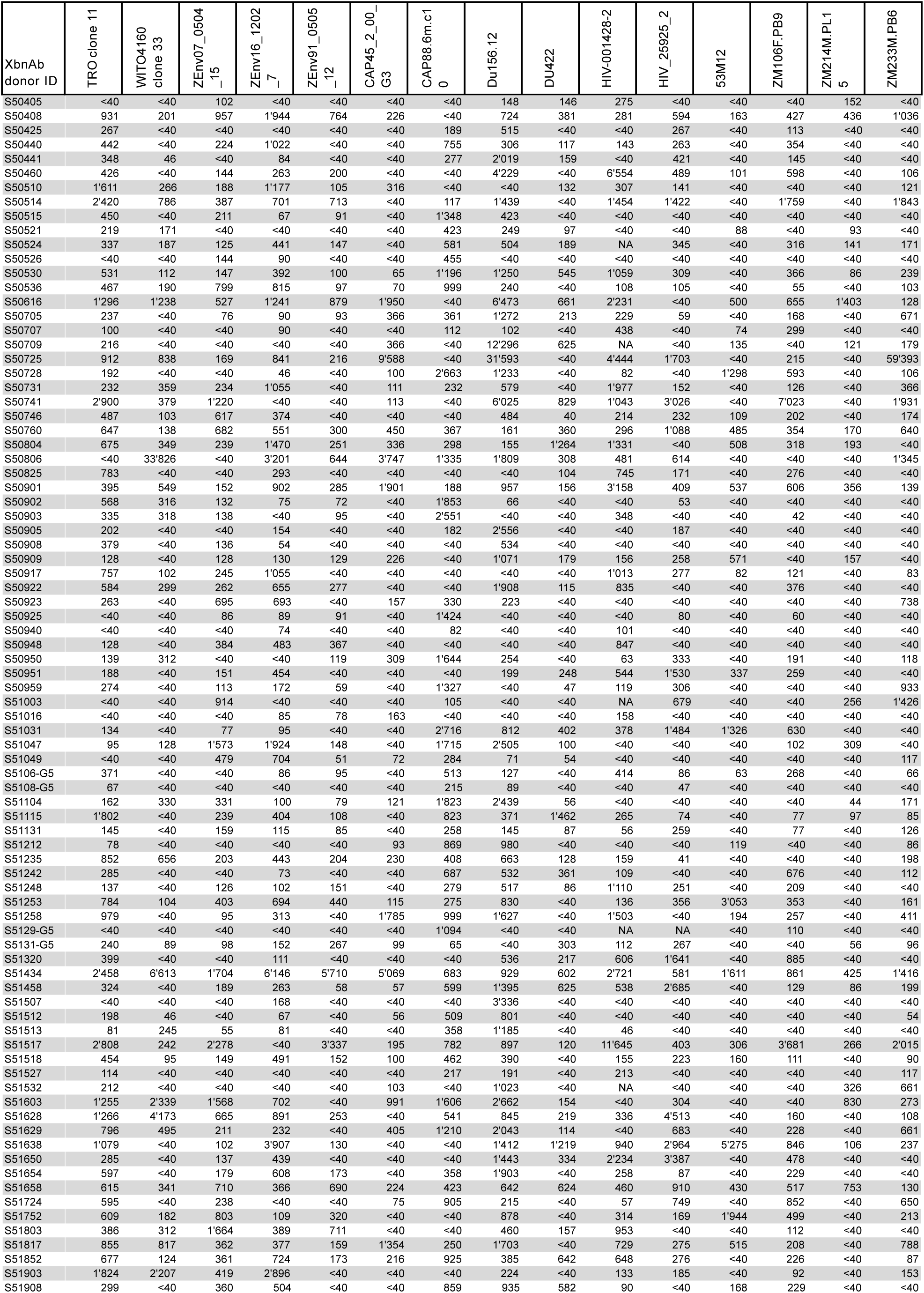
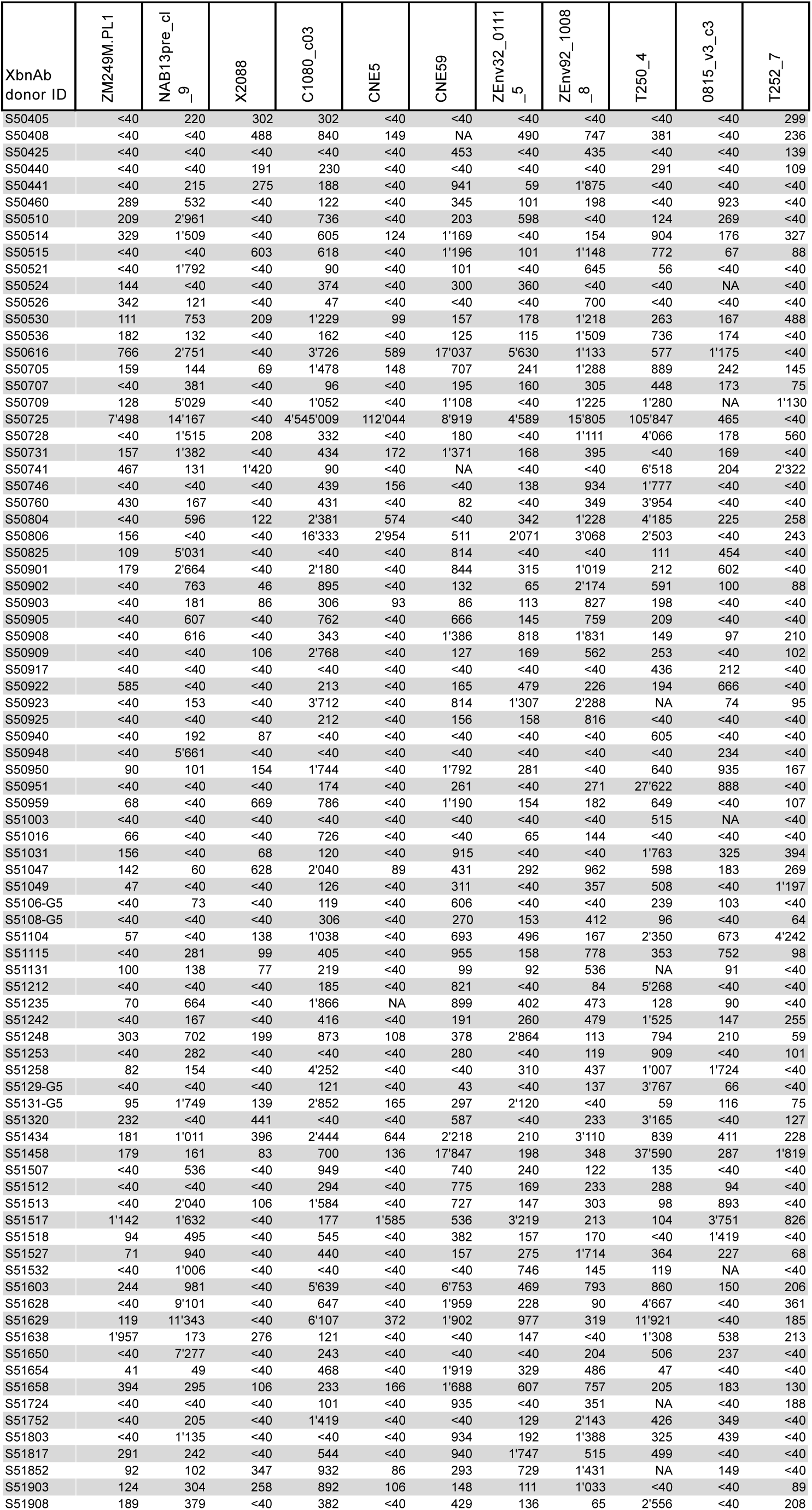
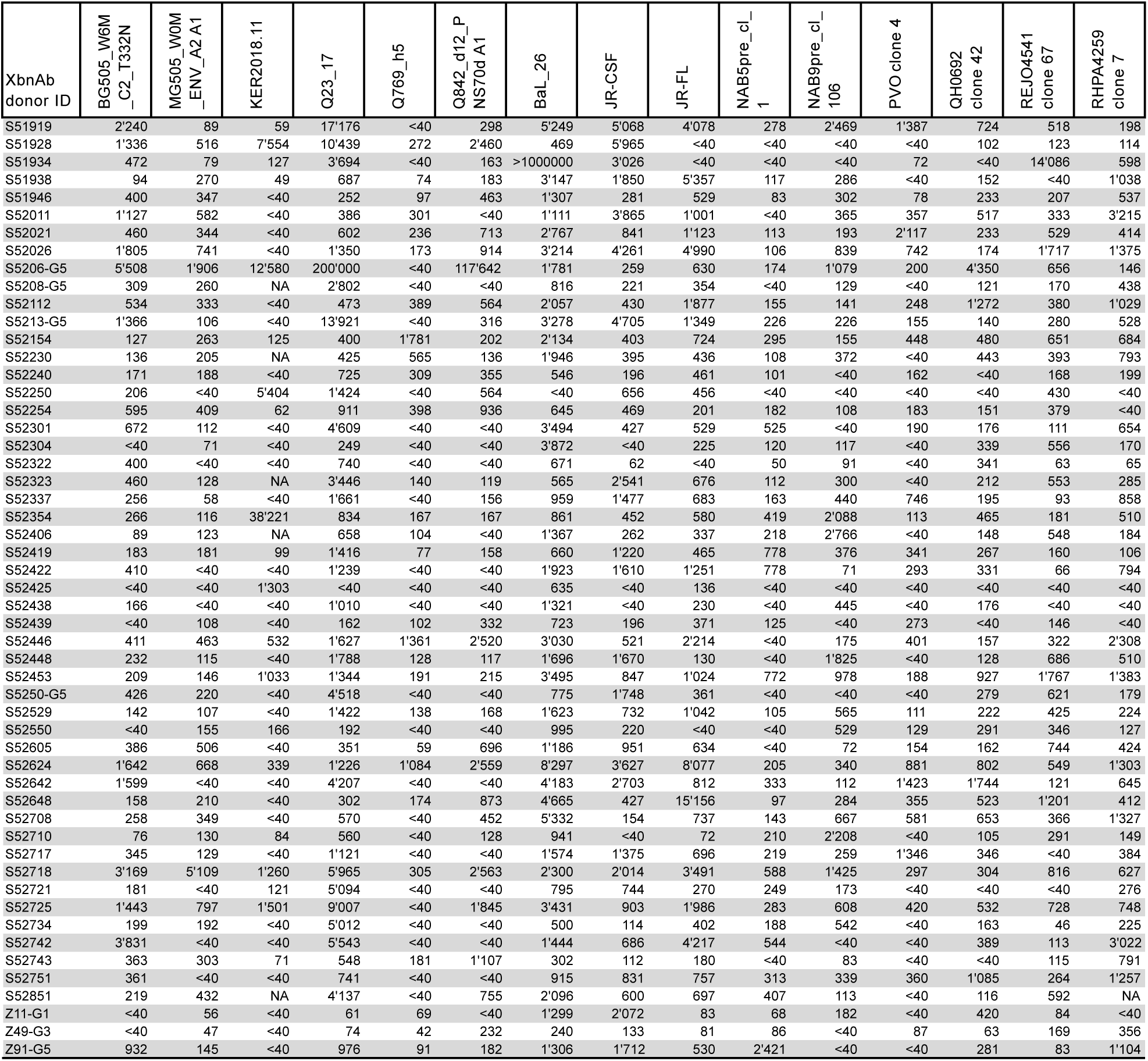
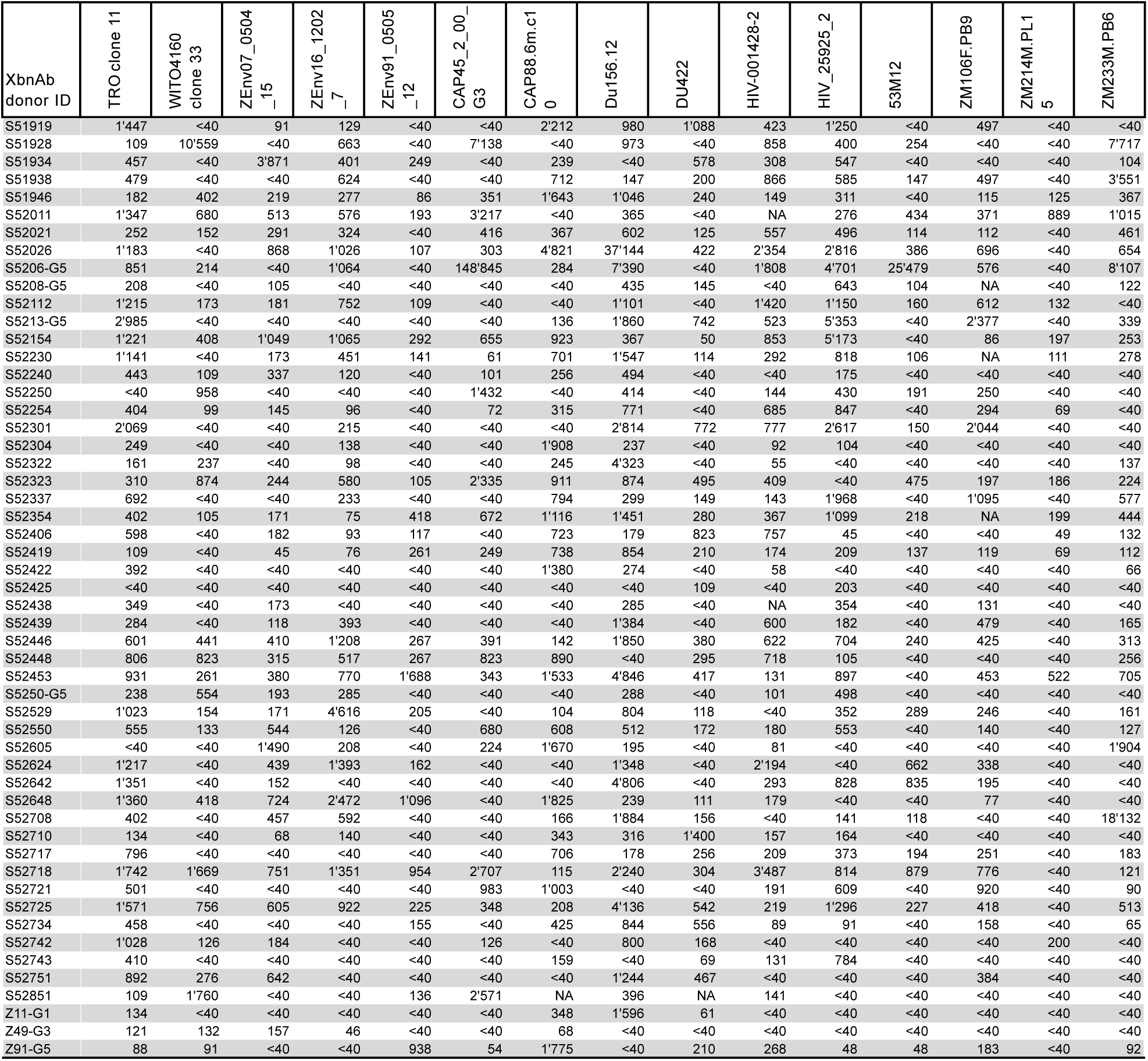
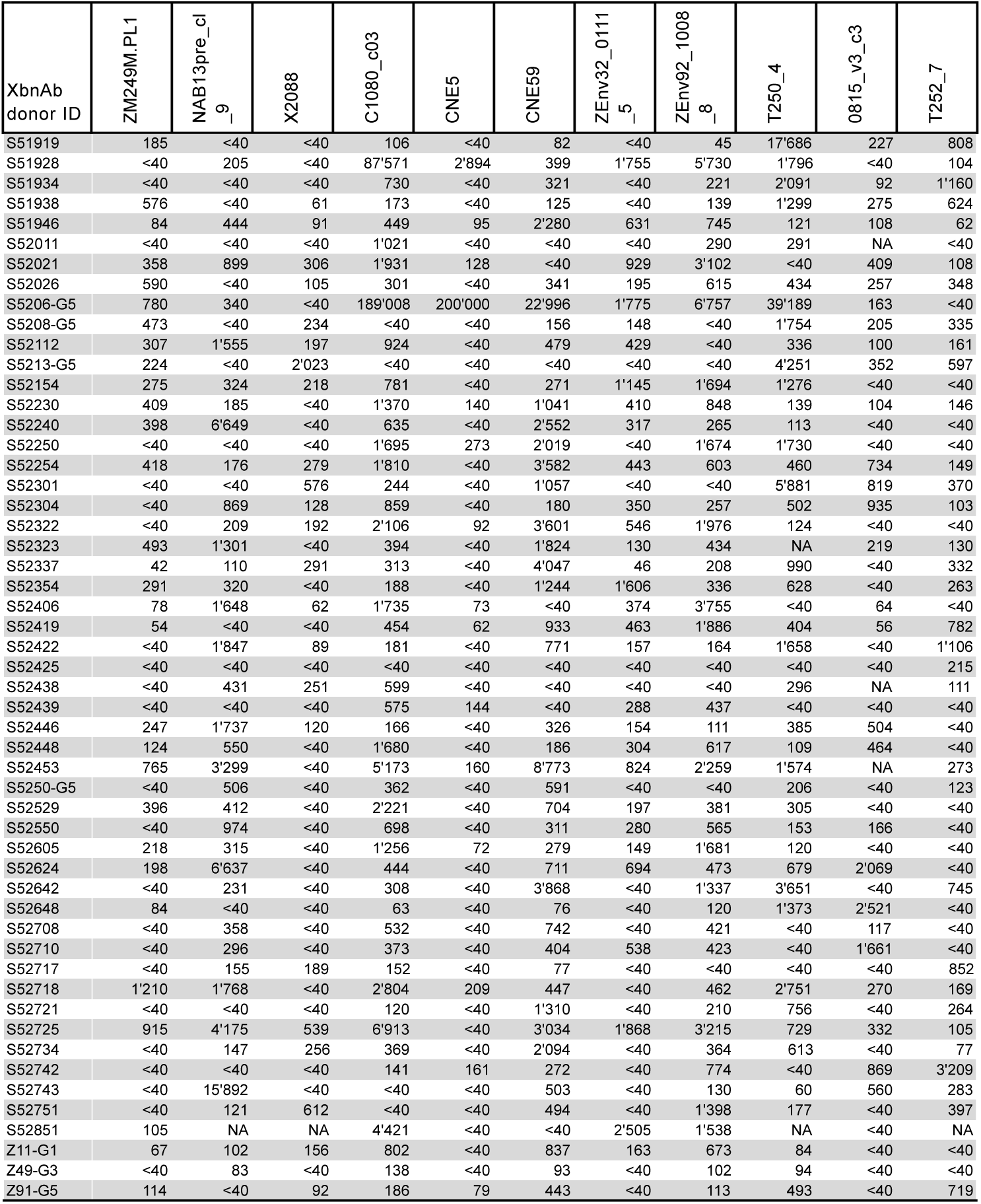
50% Neutralization titers (NT50) of XbnAb donors (n=305) against the 41-virus panel. Page 2 of 12.

**Suplementary Table 3.**
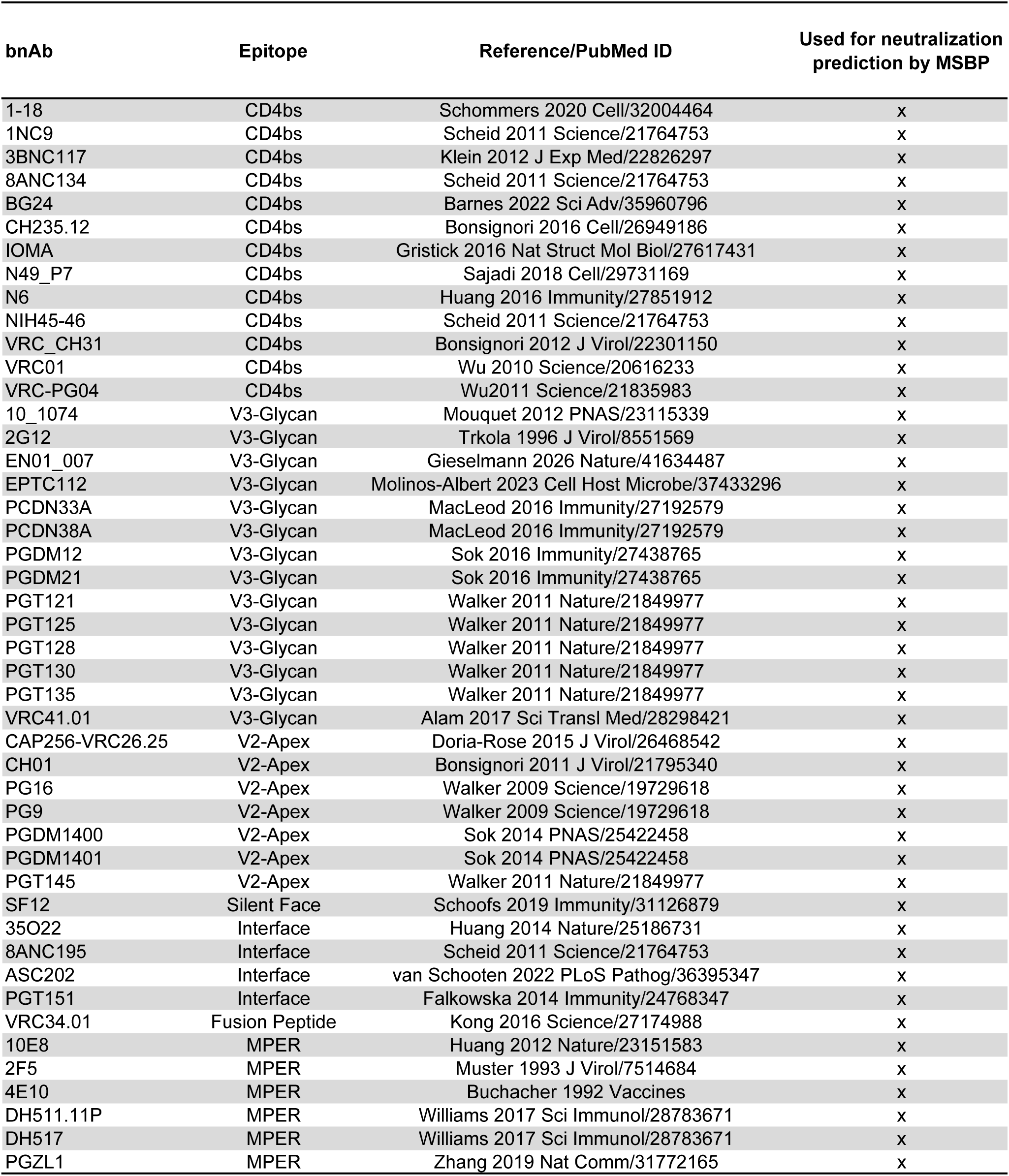
Reference bnAbs (N=46)

**Supplementary Table 4.**
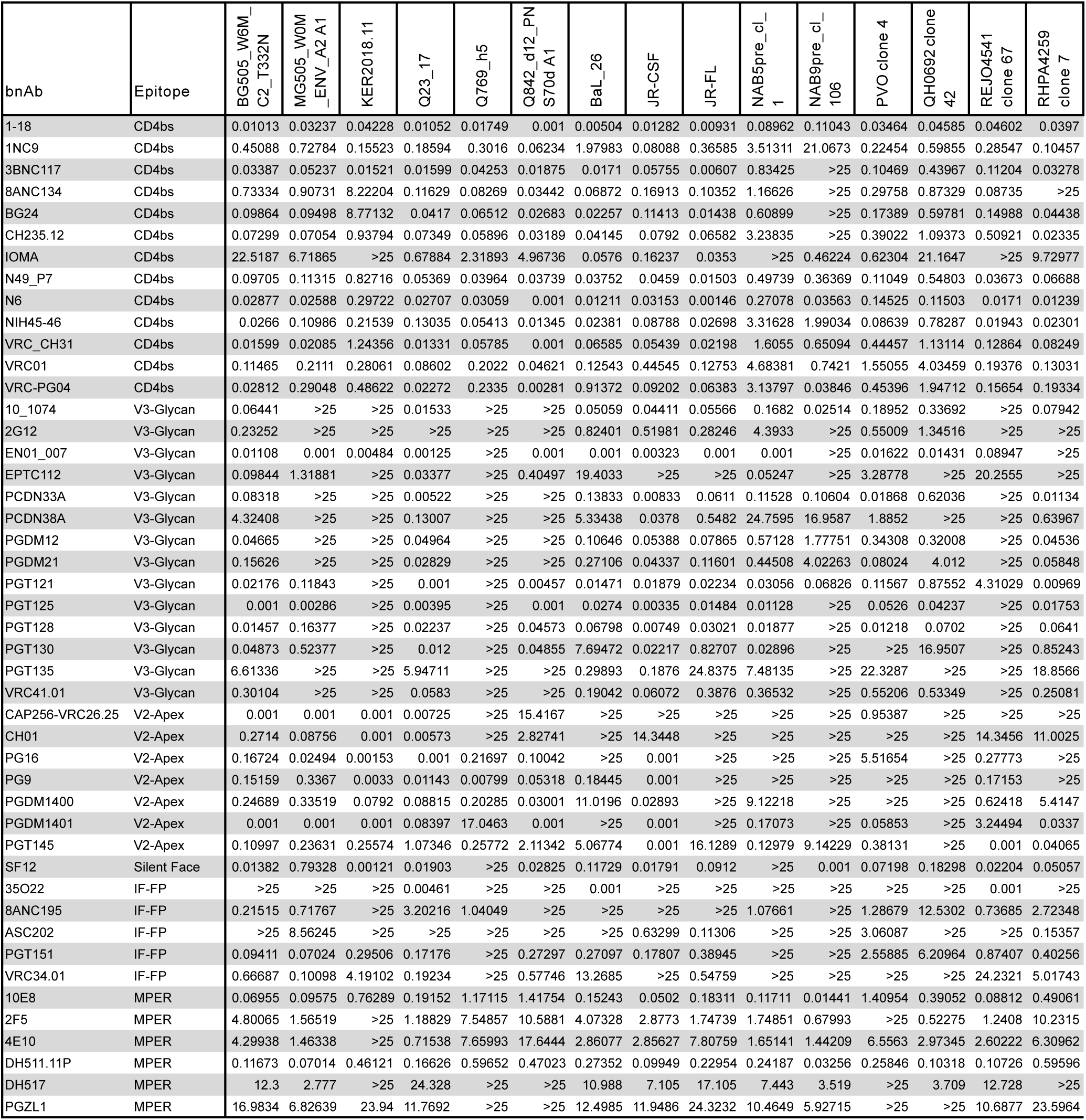
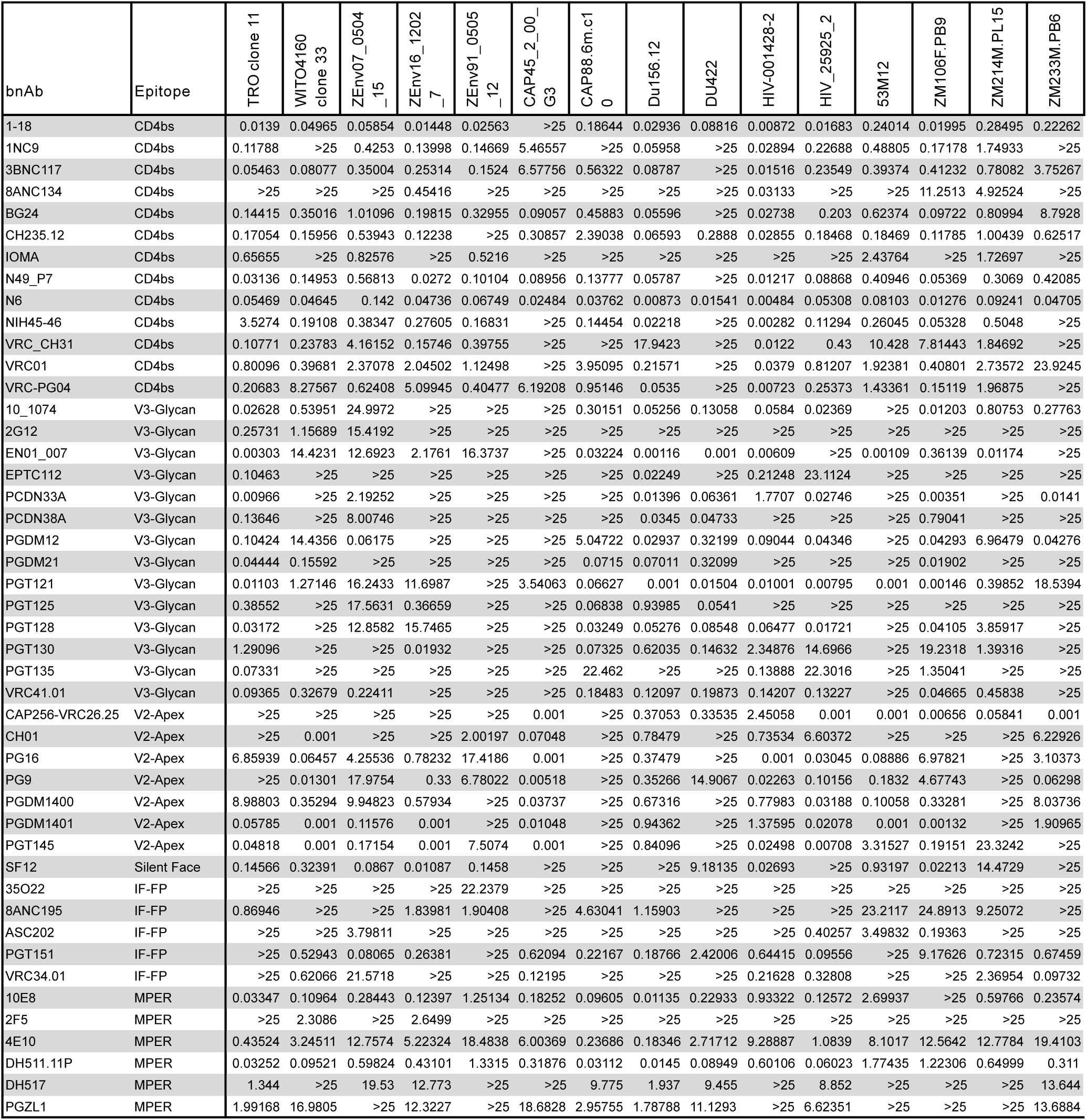
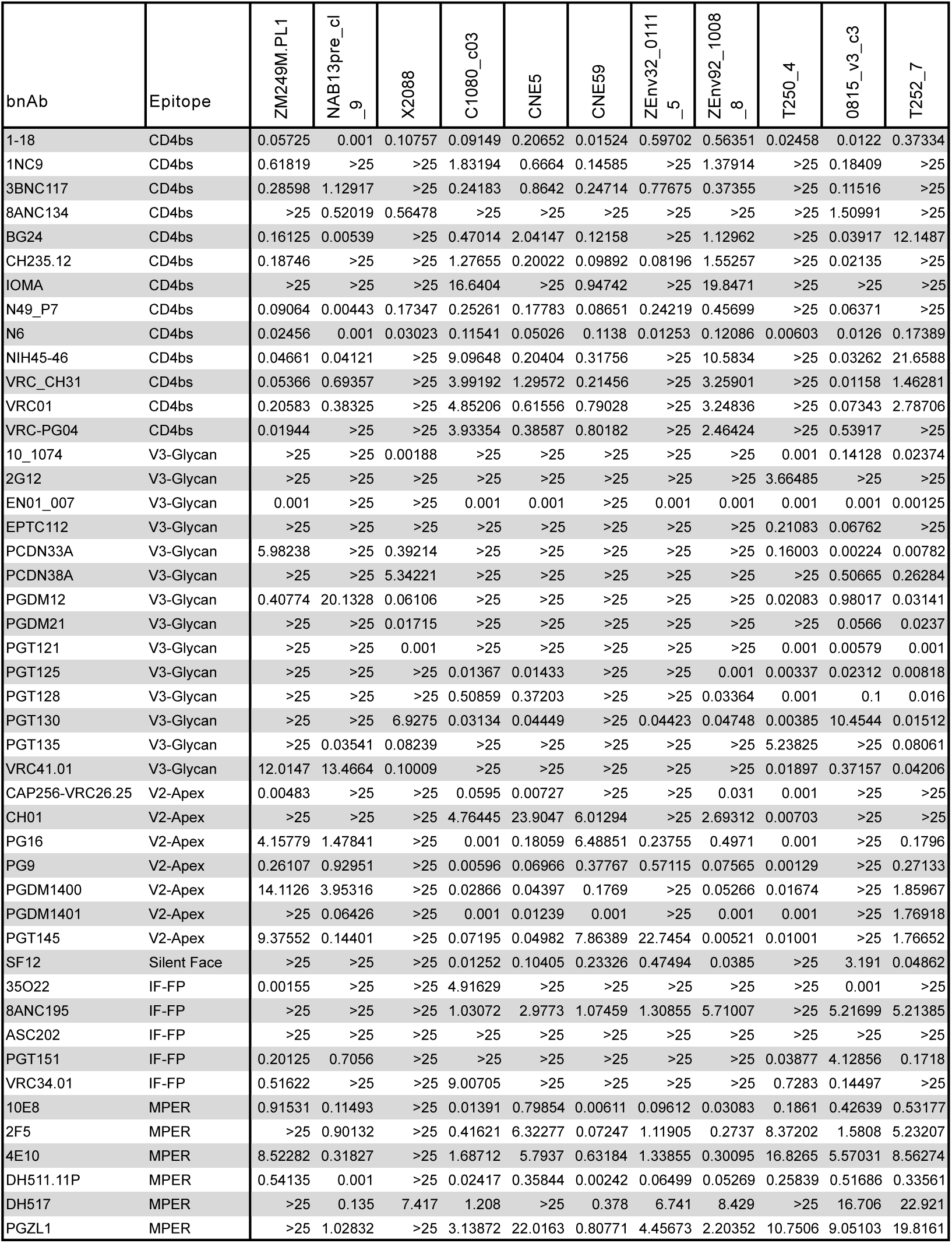
50% Inhibitory Concnetrations (IC50) of published bnAbs (n=46) against the 41-virus panel. Page 1 of 3.

